# Calcium-permeable cation channels are involved in uranium uptake in *Arabidopsis thaliana*

**DOI:** 10.1101/2021.06.10.447834

**Authors:** Manon C.M. Sarthou, Fabienne Devime, Célia Baggio, Sylvie Figuet, Claude Alban, Jacques Bourguignon, Stéphane Ravanel

**Affiliations:** Univ. Grenoble Alpes, INRAE, CEA, CNRS, IRIG, LPCV, 38000 Grenoble, France

**Keywords:** Radionuclide, higher plants, root, accumulation, transport, competition, inhibition, mutants

## Abstract

Uranium (U) is a non-essential and toxic element that is taken up by plants from the environment. The assimilation pathway of U is still unknown in plants and any other organism. In this study, we provide several evidences that U is taken up by the roots of *Arabidopsis thaliana* through Ca^2+^-permeable cation channels. First, we showed that deprivation of Arabidopsis plants with calcium induced a 1.5-fold increase in the capacity of roots to accumulate U, suggesting that calcium deficiency promoted the radionuclide import pathway. Second, we showed that external calcium inhibits U accumulation in roots, suggesting a common route for the uptake of both cations. Third, we found that gadolinium, nifedipine and verapamil inhibit the absorption of U, suggesting that different types of Ca^2+^-permeable channels serve as a route for U uptake. Last, we showed that U bioaccumulation in Arabidopsis mutants deficient for the Ca^2+^-permeable channels MCA1 and ANN1 was decreased by 40%. This suggests that MCA1 and ANN1 contribute to the absorption of U in different zones and cell layers of the root. Together, our results describe for the first time the involvement of Ca^2+^-permeable cation channels in the cellular uptake of U.

## Introduction

Trace metal elements and radionuclides are important sources of environmental pollution and hazards to human health. Uranium (U) is the heaviest naturally occurring radionuclide; its average content in Earth’s ecosystems (rocks, soils, sea and soft waters) is below 4 ppm. The main sources of anthropogenic U contamination of the environment are related to the nuclear industry and agricultural practices (Vandenhove, 2002). The fertilization of agricultural soils with phosphate fertilizers is the main source of U dispersion (Wetterlind et al., 2012). Uranium is not an essential nutrient for life but it is readily taken up by plants from the environment and can contaminate the food chain, becoming a threat to public health (Anke et al., 2009).

The most abundant U isotope (^238^U, 99.27%) is poorly radiotoxic but is highly chemotoxic to all living organisms (Gao et al., 2019). During the last decade, significant progress has been made in understanding the toxicity of U in plants (reviewed in (Chen et al., 2021). Uranium has been shown to interfere with several aspects of plant physiology, development and metabolism (Misson et al., 2009; Saenen et al., 2013; Aranjuelo et al., 2014; Vanhoudt et al., 2014; Serre et al., 2019). Biochemical, molecular and metabolomic studies indicated that U triggers oxidative stress-related response pathways and perturbs hormones synthesis and signaling, primary metabolism, iron and phosphate assimilation pathways, and cell wall synthesis (Vanhoudt et al., 2011c; Vanhoudt et al., 2011a; Doustaly et al., 2014; Saenen et al., 2015; Tewari et al., 2015; Lai et al., 2020; Lai et al., 2021). Also, U has a significant impact on mineral nutrition, with important changes in some micro- and macronutrients concentration and distribution between roots and shoots (Misson et al., 2009; Vanhoudt et al., 2011b; Doustaly et al., 2014; Berthet et al., 2018; Lai et al., 2020; Sarthou et al., 2020; Lai et al., 2021; Rajabi et al., 2021).

The uptake of non-essential elements is mediated by the assimilation pathways of essential nutrients, thanks to the broad substrate specificity of some membrane carriers and to similarities between the atomic structures of some elements. Uranium uptake and accumulation by plants is markedly influenced by its speciation in the environment. Uranium is mainly in the U(VI) form in soil solutions and its speciation is influenced by the pH, redox potential, and presence of chelating agents (e.g. carbonate, phosphate) (Chen et al., 2021). The free uranyl species (UO_2_^2+^) accumulates in roots but is poorly translocated to shoots, whereas complexation with carbonate or organic acids (e.g. citrate) triggers U translocation to shoot organs (Ebbs et al., 1998; Laurette et al., 2012a; Laurette et al., 2012b). Complexation with phosphate considerably reduces U bioavailability (by precipitation) and limits accumulation in all plant organs. In carbonate water, calcium (Ca) has been shown to modify U bioavailability and to facilitate U(VI) symplastic transport and translocation to shoots (El Hayek et al., 2018; El Hayek et al., 2019). Thus, depending on U speciation in soils, several assimilation pathways are anticipated for free or complexed anionic and cationic U species.

In spite of several studies, including transcriptomic analyses of plant responses to U stress (Doustaly et al., 2014; Lai et al., 2020; Lai et al., 2021), the assimilation pathways of the radionuclide are still not known. It has been postulated that the iron (Fe) uptake machinery could be involved in the assimilation of U since the radionuclide interferes significantly with Fe homeostasis (Doustaly et al., 2014). However, in conditions favoring free uranyl and hydroxylated species, the main Fe(II) transporter at the plasma membrane, IRT1, is not involved in U uptake (Berthet et al., 2018). The Ca assimilation pathway has been also proposed to contribute to the uptake of U by tobacco BY-2 cells, in which U bioassociation is inhibited by the cation- and Ca^2+^-channel blocker gadolinium (Rajabi et al., 2021). Moreover, Ca and magnesium (Mg), but not potassium (K), inhibit uranyl uptake with a similar efficiency in the green microalga *Chlamydomonas reinhardtii*, suggesting that Ca and Mg assimilation routes could serve for U absorption (Fortin et al., 2007).

In order to identify U assimilation pathways in land plants, we used an unbiased approach investigating the influence of nutrient deficiency on the capacity of *Arabidopsis thaliana* to accumulate U in roots. Our working hypothesis relies on the knowledge that perturbations of mineral nutrition, particularly starvations, are associated with changes in the array of plasma membrane carriers involved in mineral assimilation. For example, many ion transporters and channels are known to be upregulated under sulfate, nitrogen, iron, manganese, magnesium or potassium deficiencies (Curie et al., 2000; Vert et al., 2002; Yoshimoto et al., 2002; Kiba et al., 2012; Mao et al., 2014; Castaings et al., 2016; Lara et al., 2020). We anticipated that starvation conditions promoting U uptake could identify the routes use by U to enter root cells. Indeed, in conditions where free uranyl and hydroxylated species are predominant (acidic pH, absence of chelators), we showed that deprivation of Arabidopsis plants with Ca is mandatory for an increased bioaccumulation of U in roots, suggesting that Ca deficiency promotes the radionuclide import pathway. Then, competition and inhibitor assays, together with the analysis of Arabidopsis mutants impaired in Ca transport indicated, for the first time, that Ca^2+^-permeable cation channels are involved in U uptake in plants.

## Materials and methods

### Plant growth conditions and uranium treatment

The wild-type *Arabidopsis thaliana* ecotype Columbia (Col-0) was used in this study. Seeds were sterilized with chlorine gas (Lindsey et al., 2017) and stored in distilled water for 4 days at 4°C in the dark. Stratified seeds were sown into homemade plates thermoprinted using the amorphous polymer acrylonitrile butadiene styrene using an Ultimaker 2 3D-printer (Ultimaker BV, The Netherlands). Conical holes of the plates were filled with 0.65% (w/v) agar (Agar plant type A, Sigma-Aldrich) solubilized in distilled water. Plants were grown in hydropony in a controlled environment with an 8 h light period at 22°C (light intensity of 110 μmol of photons m^−2^s^−1^) followed by a 16 h dark period at 20°C. The nutrient solution, hereafter referred to as ‘Gre medium’, was composed of 0.88 mM K_2_SO_4_, 1 mM Ca(NO_3_)_2_, 1 mM MgSO_4_, 0.25 mM KH_2_PO_4_, 10 μM H_3_BO_3_, 0.1 μM CuSO_4_, 2 μM MnSO_4_, 0.01 μM (NH_4_)_6_Mo_7_O_24_, 2 μM ZnSO_4_, 10 μM NaCl, 0.02 μM CoCl_2_, 20 μM Fe-EDTA, and 0.25 mM MES, pH 5.6 (Table S1). Plants were grown for 29 to 33 days in Gre medium with weekly solution changes until treatment. For U treatment, plants were transferred to Gre media depleted with phosphate (100-fold dilution as compared with Gre, i.e. 2.5 μm KH_2_PO_4_) and supplemented with 20 μM uranyl nitrate (UO_2_(NO_3_)_2_). After U treatment, plants were harvested and rinsed once with 10 mM Na_2_CO_3_, then twice with distilled water to remove U that is loosely adsorbed on the root surface (Doustaly et al., 2014). Roots and shoots were then separated, dried on absorbent paper and fresh biomass was measured. Finally, roots and shoots were dehydrated at 80°C for 24 h and dry weighed.

A similar procedure was used to analyze U bioaccumulation in *A. thaliana* mutant lines deficient in Ca^2+^-permeable cation channels. The *mca1*-null, *mca2*-null, and the double *mca1/mca2*-null mutants were kindly provided by Prof. Iida and Prof. Miura (Nakagawa et al., 2007; Yamanaka et al., 2010). Seeds of an *ann1*-null mutant (Lee et al., 2004) were obtained from the European Nottingham Arabidopsis Stock Centre (N661601, SALK_015426C).

### Mineral depletion assays

After 29 days of growth in standard Gre medium, plants were transferred for 4 days in a modified medium in which one or several nutrients were depleted by a 100-fold as compared with the standard concentrations (Table S1). After 4 days of starvation, plants were transferred into the same modified Gre medium in which phosphate was lowered to 2.5 μM and uranyl nitrate was added at 20 μM. Plants were cultivated for 24 h in these conditions and then harvested. Photosystem II efficiency in dark-adapted leaves (Fv/Fm) was measured using a FluorPen (FP100/D, Photon Systems Instruments, Brno, Czech Republic) and used as a proxy for plant fitness in mineral-depleted media.

### Competition assays

After 33 days of growth in standard Gre medium, plants were rinsed with distilled water and transferred into a solution containing 0.25 mM MES (pH 5.6), 20 μM uranyl nitrate and various concentrations (20, 200 or 2000 μM) of Ca(NO_3_)_2_. Plants were harvested after 10 h of treatment in these solutions, at 22°C in the light (110 μmol of photons m^−2^s^−1^). Competition assays were also performed with KNO_3_ or K_2_SO_4_ in a similar concentration range.

### Inhibitors assays

After 33 days of growth in standard Gre medium, plants were rinsed with distilled water and transferred in 0.25 mM MES (pH 5.6) containing either 250 μM gadolinium chloride (GdCl_3_), 100 μM verapamil hydrochloride, 50 μM nifedipine or 1% (v/v) DMSO for 20 min at 22°C. Uranyl nitrate 20 μM was then added to different solutions and plants were harvested 2 or 6 h after U addition (22°C at 110 μmol of photons m^−2^s^−1^). Verapamil and nifedipine stock solutions (100-fold concentrated) were prepared in DMSO; gadolinium chloride was solubilized in distilled water.

### Inductively coupled plasma mass spectrometry (ICPMS) analyses

Dried samples were mineralized in 400 μL of 65% (w/v) HNO_3_ (Suprapur, Merck) for 3 h using a HotBlock CAL3300 (Environmental Express) digestion system. Digested samples were diluted in distilled water at 0.65% (w/v) HNO_3_ and analyzed using an iCAP RQ quadrupole mass instrument (Thermo Fisher Scientific GmbH, Germany). Elements (^24^Mg, ^25^Mg, ^31^P, ^39^K, ^43^Ca, ^44^Ca, ^55^Mn, ^56^Fe, ^57^Fe, ^64^Zn, ^66^Zn, ^95^Mo, ^98^Mo, ^238^U) were analyzed using the collision mode with helium as a cell gas. Elements concentrations were determined using standard curves and corrected using an internal standard solution of ^45^Sc, ^103^Rh, ^172^Yb.

### Predictions of uranium speciation

Visual MINTEQ (version 3.1, https://vminteq.lwr.kth.se/) was used to predict U speciation in the different media. The pH and temperature parameters were set at 5.6 and 20°C, respectively. The concentrations of aqueous inorganic species (in mol/L) and the distribution among dissolved and soluble species (in %) were obtained using default settings.

### Statistical analyses

Statistical analyses were performed using the R Studio software (RStudio Team, 2015) and the *nparcomp* package (Konietschke et al., 2015). Non-parametric statistical analysis was done on our datasets that typically contain small sample sizes (n=4 plants in each assay) and do not meet the assumptions of parametric tests (normal distribution and homogeneity of variance, as determined using the Shapiro–Wilk and Fisher tests, respectively). Non-parametric Tukey tests were conducted and the confidence level was set at 99% (*p*<0.01).

## Results

### Multi-element mineral deprivations improve U accumulation in roots

To evaluate the impact of nutrient deprivations on the accumulation of U in *Arabidopsis thaliana*, plants were grown for 4 weeks in a complete standard Gre medium and then transferred for 4 days in a Gre medium depleted of all nutrients (100-fold dilution of Gre, Gre/100 medium) (Table S1). After 4 days of culture, nutrient-deprived plants displayed shorter roots and smaller, darker green leaves as compared with control plants. The photosynthetic parameter Fv/Fm (photosystem II efficiency in dark-adapted leaves) was used as a proxy to evaluate plant fitness. Fv/Fm of plants grown in Gre/100 was not changed as compared with plants maintained in Gre (0.79±0.02 in both conditions). Roots and shoots were digested with nitric acid and their ionome was analyzed by ICPMS. Nutrient deprivation had a strong impact on root and shoot ionomes, with a significant decrease in the pools of Mg, P, K, Ca, Mn, Fe, Zn and Mo (Figures S1, S2; Tables S2, S3). For U bioaccumulation assays, control and deprived plants were challenged with 20 μM uranyl nitrate for 24 h in Gre/100 or Gre medium in which the phosphate concentration was reduced by 100-fold to limit the precipitation of U and preserve its bioavailability (Ebbs et al., 1998; Misson et al., 2009; Laurette et al., 2012b). Plants were extensively washed with sodium carbonate and distilled water to remove loosely-bound U and the root and shoot ionomes were analyzed by ICPMS. Nutrient-deficient plants displayed a 3-fold increase of U associated with roots as compared with control plants (Figure 1). The accumulation of U in shoots was similar in the two conditions (data not shown). Uranium is characterized by a low root-to-shoot translocation rate in the absence of ligands (e.g. carbonate, organic acids) in the environment (Ebbs et al., 1998; Laurette et al., 2012b). In our experimental conditions the translocation factor was in the range 2-7×10^−4^, meaning that less than 0.1% of U accumulated in roots was translocated to shoots. Due to the very low translocation rate, the U content in shoots will not be discussed in the rest of the study.

**Figure 1:**
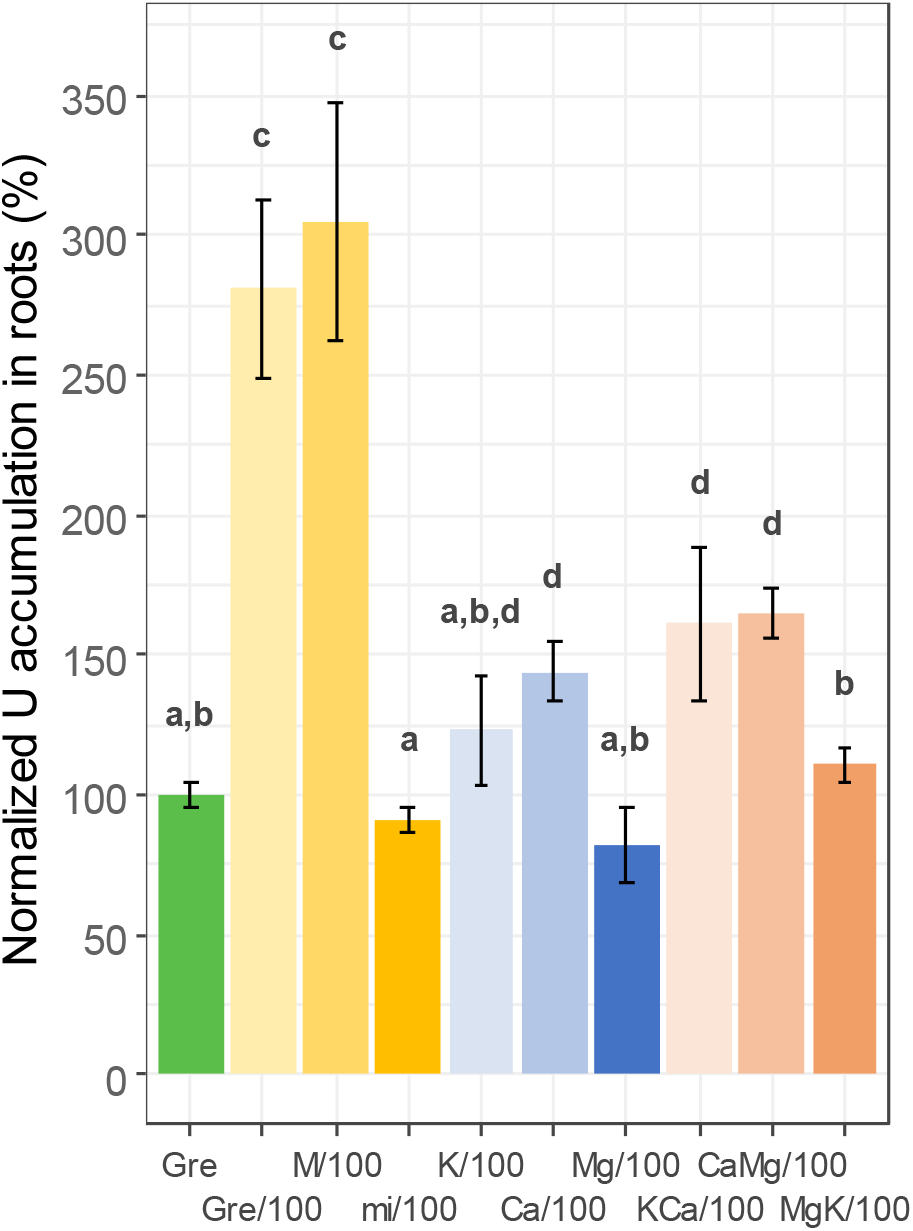
Bioaccumulation of U in the roots of Arabidopsis plants deprived with nutrients. Four-week-old Arabidopsis plants grown in a complete standard Gre medium were transferred for 4 days in Gre medium depleted of one or several nutrients (100-fold reduction of the standard concentration; Table S1). Plants were deprived with all elements (Gre/100), macroelements (M/100), microelements (mi/100), or Ca, K, and Mg, alone or in dual combination. Nutrient-deprived plants were challenged with 20 μM uranyl nitrate for 24 h, thoroughly washed with sodium carbonate and distilled water to remove loosely-bound metals, and U was measured by ICPMS in mineralized roots. Data from 2 to 11 replicates of the experiment have been gathered, with n=4 plants for each condition. To normalize fluctuations between replicates, U accumulated in roots was expressed as a percentage related to the control condition in Gre medium. Bar plots represent mean ± SD. Letters indicate significant differences determined using a non-parametric Tukey’s test with p<0.01. The amount of U bioaccumulated in Gre medium was 727 ± 79 μg.g^−1^ fresh weight (mean ± SD, n=11 independent experiments).

In order to identify which changes in nutrient supply were responsible for the increase of U associated with roots in Gre/100, 4-week-old Arabidopsis plants were starved with either micronutrients (B, Co, Cu, Fe, Mn, Mo and Zn; mi/100 medium) or macronutrients (Ca, K, Mg, P, S and N; M/100 medium). After 4 days of starvation, the roots and shoots of plants grown in mi/100 were significantly depleted in Fe, Mn, Mo and Zn (B, Co, and Cu were not quantified) (Figures S1, S2; Tables S2, S3) but had no distinguishable phenotype as compared with plants grown in Gre. Plants grown in macronutrient-deprived medium had reduced pools of Mg, P, K and Ca (S and N were not quantified) (Figures S1, S2; Tables S2, S3). As compared with controls, M/100 plants had shorter and more branched roots, smaller and greener shoots, but comparable Fv/Fm values (0.79±0.02). After 24 h of U treatment, no significant change of U content was measured in roots of micronutrient-deficient plants as compared with Gre. However, macronutrient-deficient plants showed a 3-fold increase of U associated with roots as compared with control (Figure 1). U content in roots from M/100 plants was similar to that of complete nutrient-deprived plants (Gre/100).

Noteworthy, the 100-fold reduction in the cations Ca^2+^, Mg^2+^ and K^+^ in M/100 medium is accompanied by important changes in the concentration of the counter-anions NO_3_^−^, SO_4_^2-^ and PO_4_^3-^ (100-, 82- and 100-fold reduction, respectively) (Table S1). To analyze the effect of sulfate starvation on the accumulation of U in roots, (NH_4_)_2_SO_4_ was added to M/100 to recover the concentration of the control Gre medium. Root accumulation of U in M/100 supplemented with (NH_4_)_2_SO_4_ was similar to the one in M/100 plants (Figure S3). To test the effect of phosphate starvation, plants were deprived with phosphate (100-fold reduction; P/100) for 4 days and challenged with 20 μM uranyl nitrate for 24 h. In this condition, plants displayed no phenotype and accumulated similar amount of U in roots as compared with plants in the control Gre medium (Figure S3). The impact of nitrate depletion was addressed in the M/100 medium supplemented with either KNO_3_ or Mg(NO_3_)_2_. These conditions correspond to double cation deprivations (Ca and Mg or Ca and K) and will be discussed below; they showed that nitrate depletion has no impact on the amplitude of U bioaccumulation in roots (Figure S3).

### Uranium speciation is not significantly modified in nutrient-depleted media and do not influence U bioaccumulation in roots

At this stage, changes in U content in roots in Gre/100 and M/100 conditions could be due to an increased capacity of roots to accumulate U because of modifications induced by the triple Ca-Mg-K deprivation, and/or to changes in U speciation and bioavailability in the different media. The second hypothesis was addressed by modelling U speciation using the Visual MINTEQ software. The chemical forms of U that are predicted in the control Gre medium (pH 5.6) are mainly cationic, with about 70% of hydroxylated species and 9% of free UO_2_^2+^ (Table S1). In M/100, these proportions are slightly changed (79% of hydroxylated forms, 7% of free UO_2_^2+^) but cannot explain the 3-fold increase in U accumulation in the roots of M/100 plants as compared with Gre plants. The main change was observed for the UO_2_SO_4_ form, which represents about 8% of U species in Gre and 0.2% in M/100 (Table S1). Supplementation of M/100 with (NH_4_)_2_SO_4_ restored the proportion of UO_2_SO_4_ species at a level (6%) comparable to that of the Gre medium and did not change the high accumulation of U in roots (Figure S3). Together, these predictions indicated that the speciation of U has no or limited role in the different capacity of roots to accumulate U in our experimental conditions.

### Calcium deprivation is essential for an increased accumulation of U in roots

To assess which of the 3 elements (Ca, Mg and K) is important for the improvement of U accumulation in nutrient-starved Arabidopsis roots, bioaccumulation assays were done using single-nutrient deprived Gre media (Ca/100, Mg/100 or K/100). Mg/100 plants were strongly deficient in Mg in both roots and shoots (40% decrease as compared with controls) (Figures S1, S2; Tables S2, S3) but did not show a significant change in U accumulation in roots relative to Gre (Figure 1). K/100 roots showed a moderate (<10%) decrease in K content (Figure S1; Table S2) and did not accumulate more U than control plants in Gre (Figure 1). Calcium deficiency was important in Ca/100 plants, with a 2-fold reduction of the root and shoot Ca pools (Figures S1, S2; Tables S2, S3). A 50% increase of the accumulation of U in the roots of Ca/100 plants was measured as compared with Gre plants (Figure 1).

The increase of U accumulation in the roots of plants deprived with Ca alone was not as important as in M/100 or Gre/100 plants. Thus, we tested the effect of double nutrient deprivations on the accumulation of U in roots. Plants starved with Mg and K (MgK/100 medium) were deficient in both cations (Figures S1, S2; Tables S2, S3) and accumulated similar amount of U in roots as control plants in Gre (Figure 1). In contrast, plants starved with K and Ca (KCa/100) or Ca and Mg (CaMg/100) were strongly deficient in Ca (Figures S1, S2; Tables S2, S3) and showed a 50% increase of U associated with roots when compared with Gre plants (Figure 1). Double-cation deprived media were also obtained using a M/100 medium in which KNO_3_ or Mg(NO_3_)_2_ was added to restore the concentration of K^+^ or Mg^2+^ to standard levels. These conditions also abolished nitrate depletion due to Ca(NO_3_)_2_ limitation (1/100 dilution). In these conditions, U accumulation levels in roots were similar to those measured in Ca/100, KCa/100 or CaMg/100 (Figure S3).

Uranium speciation in the single- and double-deprived Gre media was computed using Visual MINTEQ. As for Gre and M/100, the main chemical forms of U were predicted to be hydroxylated species (69 to 80% of total U forms) and free UO_2_^2+^ (about 8%), with little variation from one medium to the other (Table S1). Significant changes were predicted for UO_2_SO_4_, which represents about 5% of U species in K/100, Mg/100, KCa/100 and CaMg/100, 9% in Ca/100, 0.3% in MgK/100 (Table S1). There is no correlation between the proportion of the UO_2_SO_4_ species and the accumulation of U in roots, indicating again that U speciation does not modify significantly U adsorption/absorption in our experimental conditions.

Together, these data showed that a decrease of Ca content in Arabidopsis plants is necessary for an improved accumulation of U in roots. Perturbations of Mg and/or K homeostasis alone do not have similar consequences.

### Uranium adsorption in the apoplast and absorption in the symplast are influenced by changes in nutrient supply

A bioaccumulation assay was done in the cold to discriminate between U adsorption or absorption by roots, i.e. to address the distribution of the radionuclide between the apoplast and the symplast of root cells. Four-week-old plants were transferred into Gre, M/100 or Ca/100 for 4 days, pre-incubated for 1 h at either 4 or 22°C for acclimation, and challenged with U for 24 h at 4°C or 22°C under a standard light regime. Cold-treated plants still have the capacity to accumulate significant amount of U in roots in all conditions, but with a 1.4 to 1.7-fold reduction as compared with plants incubated at 22°C (Figure 2). Considering that cold-treatment strongly, but not completely, impairs ion transport processes, one can estimate that at least 30-40% of bioaccumulated U has been taken up by root cells. The rest of U is specifically adsorbed in the apoplast (roots have been thoroughly washed with sodium carbonate to remove loosely-bound elements). Noteworthy, cold-treated M/100 and Ca/100 roots still accumulated about 3-time and 1.5-time more U than the cold-treated control roots (Figure 2), respectively, meaning that the Ca-Mg-K or Ca deprivations induced changes in the capacity of U adsorption and absorption through the apoplastic and symplastic routes, respectively.

**Figure 2:**
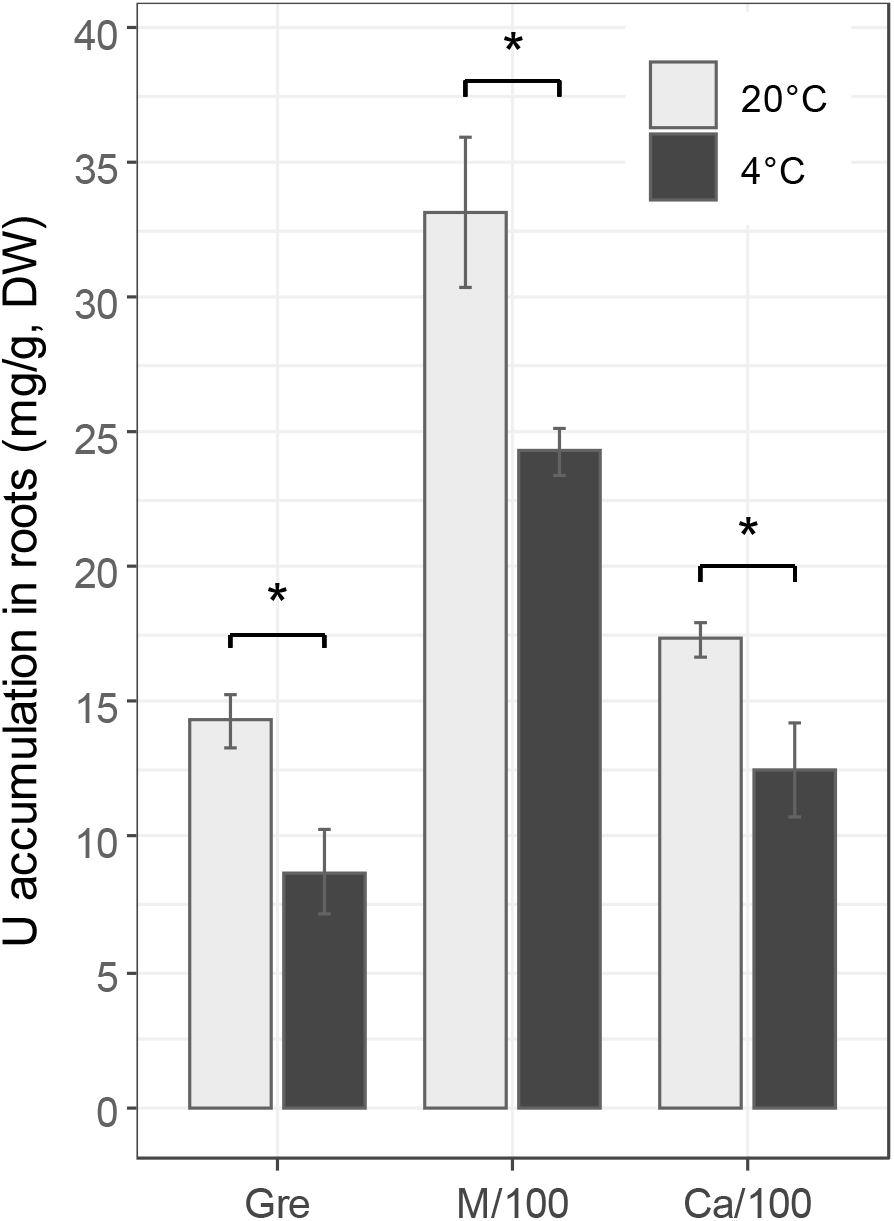
Effect of temperature on the accumulation of U in roots. Four-week-old Arabidopsis plants were transferred in Gre, M/100 or Ca/100 media for 4 days and challenged with 20 μM uranyl nitrate for 24 h at either 20 or 4°C in a standard light regime. Uranium (mg.g^−1^ dry weight) was measured by ICPMS. Bar plots represent mean ± SD with n=4 biological replicates. Stars indicate significant differences determined using a non-parametric Tukey’s test with p <0.001.

### Calcium competes with U for root uptake

Competition assays were designed to reinforce the relationship between Ca homeostasis and U accumulation by roots. Four-week-old Arabidopsis plants were challenged for 10 h with 20 μM uranyl nitrate in the presence of increasing concentrations of Ca(NO_3_)_2_ in 0.25 mM MES (pH 5.6). A very simple incubation medium was used to limit changes in U speciation and to accurately measure the effects of competition. Visual MINTEQ predictions indicated that hydroxylated and free uranyl species account for 91-93% and 7-9% of total U, respectively, in competition assays containing zero to 2 mM Ca(NO_3_)_2_ (Table S1). At equimolar concentration (20 μM), Ca had no effect on U accumulation in roots but a significant decrease was observed for 10- and 100-fold Ca excess (Figure 3). Competitions assays with similar concentrations of K_2_SO_4_ or KNO_3_ did not show any change in U accumulation (data not shown), indicating that Ca^2+^ but not K^+^ or NO_3_^−^ competes with U for root accumulation.

**Figure 3:**
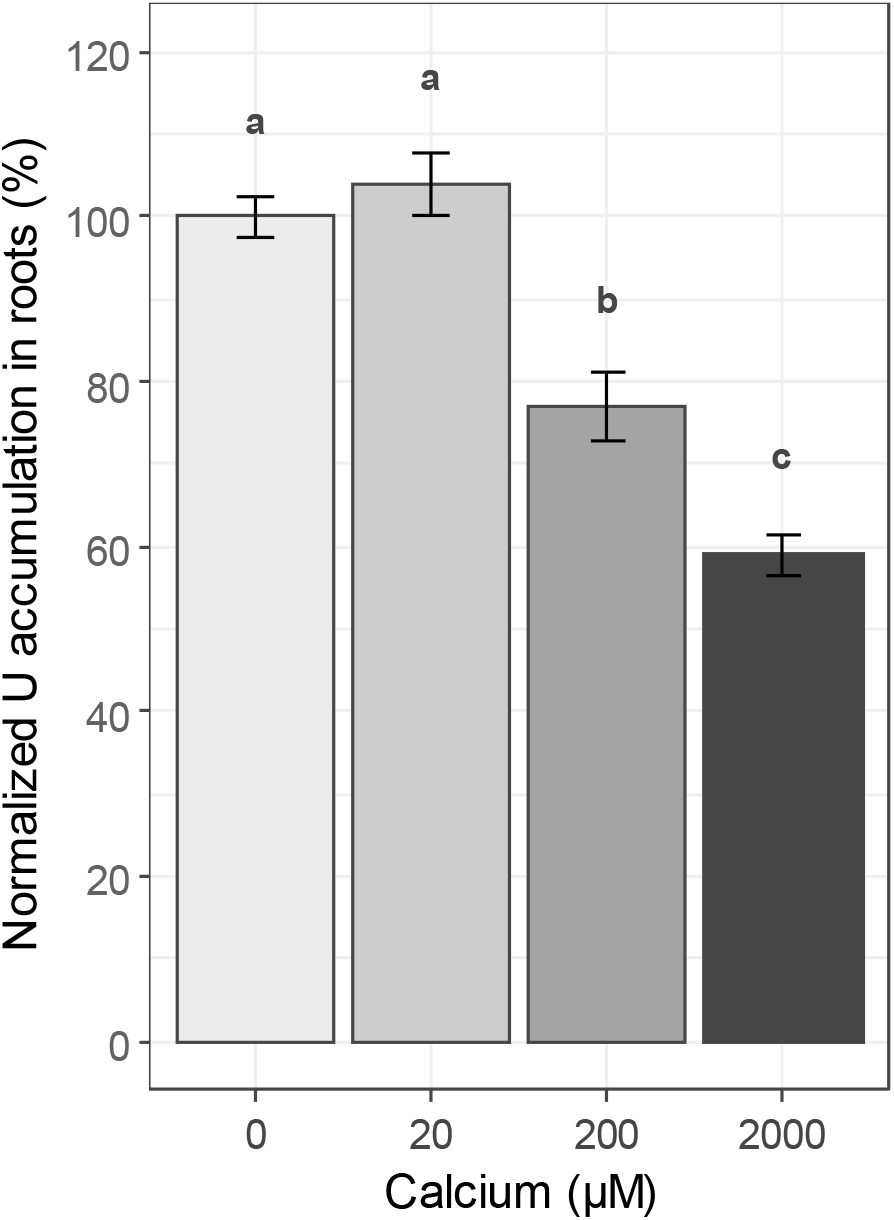
Competition between calcium and U for root uptake. Four-week-old Arabidopsis plants were challenged for 10 h with 20 μM uranyl nitrate in presence of increasing concentrations of Ca(NO_3_)_2_ in 0.25 mM MES (pH 5.6). Uranium content was measured by ICPMS and standardized relative to the control condition without calcium nitrate. The amount of U bioaccumulated in MES medium was 846 ± 79 μg.g^−1^ fresh weight (mean ± SD, n=8 measurements). Bar plots represent mean ± SD with n=4 to 8 biological replicates. Letters indicate significant differences determined using a non-parametric Tukey’s test with p <0.01.

### Uranium uptake by roots is inhibited by calcium channels blockers

A pharmacological approach was used to gain insights into U absorption by roots and to test the involvement of Ca channels. To this aim we used three compounds, gadolinium (Gd^3+^), a lanthanide used to block non-selective cation channels (NSCCs) and mechanosensitive channels (MSCCs), and verapamil and nifedipine, which are antagonists of voltage-dependent L-type Ca^2+^-permeable channels (VDCCs) (De Vriese et al., 2018). Four-week-old Arabidopsis plants were preincubated for 20 min at 22°C in 0.25 mM MES (pH 5.6) containing either 250 μM GdCl_3_, 100 μM verapamil, 50 μM nifedipine or 1% (v/v) DMSO (used as a solvent for verapamil and nifedipine). Plants were then challenged with 20 μM uranyl nitrate for 2 and 6 h at 22°C. Gadolinium inhibited U accumulation in roots by 40 to 60% throughout the kinetic (Figure 4A). Uranium speciation was not changed in Gd-containing assays as compared with controls in MES (Table S1). After 2 h of treatment, verapamil and nifedipine had little impact on U accumulation in roots but the inhibition was important (30 to 45% decrease) after 6 h of incubation (Figure 4B). In summary, the pharmacological approach suggested that some types of Ca^2+^-permeable channels, e.g. NSCCs, MSCCs and VDCCs, are involved in U uptake by roots.

**Figure 4:**
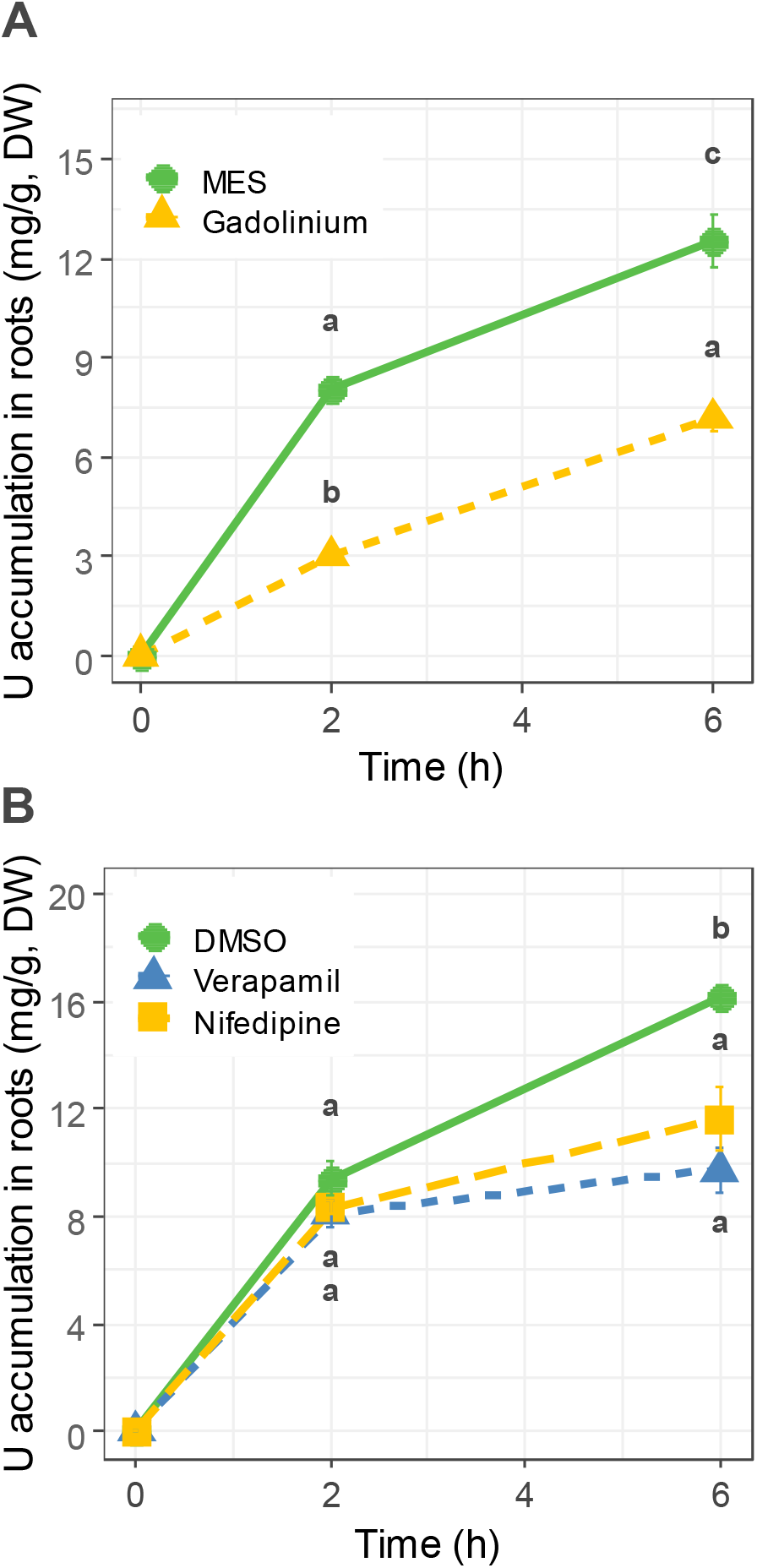
Effect of calcium channel inhibitors on the uptake of U by roots. Four-week-old Arabidopsis plants were preincubated for 20 min at 22°C in 0.25 mM MES (pH 5.6) containing (A) 250 μM GdCl_3_, or (B) 100 μM verapamil, 50 μM nifedipine or 1% (v/v) DMSO (used as a solvent for verapamil and nifedipine). Uranium (mg.g^−1^ dry weight) was measured after 2 and 6 h of incubation at 22°C in the presence of 20 μM uranyl nitrate. Curves represent mean ± SD with n=4 biological replicates. Letters indicate significant differences determined using a non-parametric Tukey’s test with p <0.01.

### Mutants in the Ca^2+^-permeable mechanosensitive MCA1 channel and in ANNEXIN1 are impaired in U uptake by roots

To gain insight into the mechanisms contributing to U uptake by roots we analyzed the accumulation of the radionuclide in Arabidopsis mutants impaired in the transport of Ca. Two Ca^2+^-permeable mechanosensitive channels localized in the plasma membrane, named MCA1 and MCA2 for *mid1*-complementing activity, have been identified in *A. thaliana* (Nakagawa et al., 2007; Yamanaka et al., 2010). Single *mca1* and *mca2* knockout mutants and the double *mca1/mca2*-null line (Yamanaka et al., 2010) were grown for 4 weeks in Gre medium and challenged with 20 μM uranyl nitrate in Gre depleted with phosphate. Short-time uptake kinetics (6 h) indicated that the *mca1* and *mca1/mca2* lines accumulated 40% less U in roots than the wild-type Col-0 (Figure 5). A small (10%) but significant decrease of U bioaccumulation was also measured in the *mca2* single mutant. Uranium bioaccumulation assays were also done with an ANNEXIN1 *ann1*-null mutant. Members of the plant annexin family function as unconventional Ca^2+^-permeable channels with roles in development and stress signaling (Davies, 2014). After 6 h of treatment with 20 μM uranyl nitrate, the *ann1*-null mutant displayed a strong reduction (40%) of U accumulation in roots as compared with wild-type plants (Figure 5). Together, these results showed that mutants in the Ca^2+^-permeable mechanosensitive MCA1 channel and in ANNEXIN1 are highly impaired in U uptake by roots.

**Figure 5:**
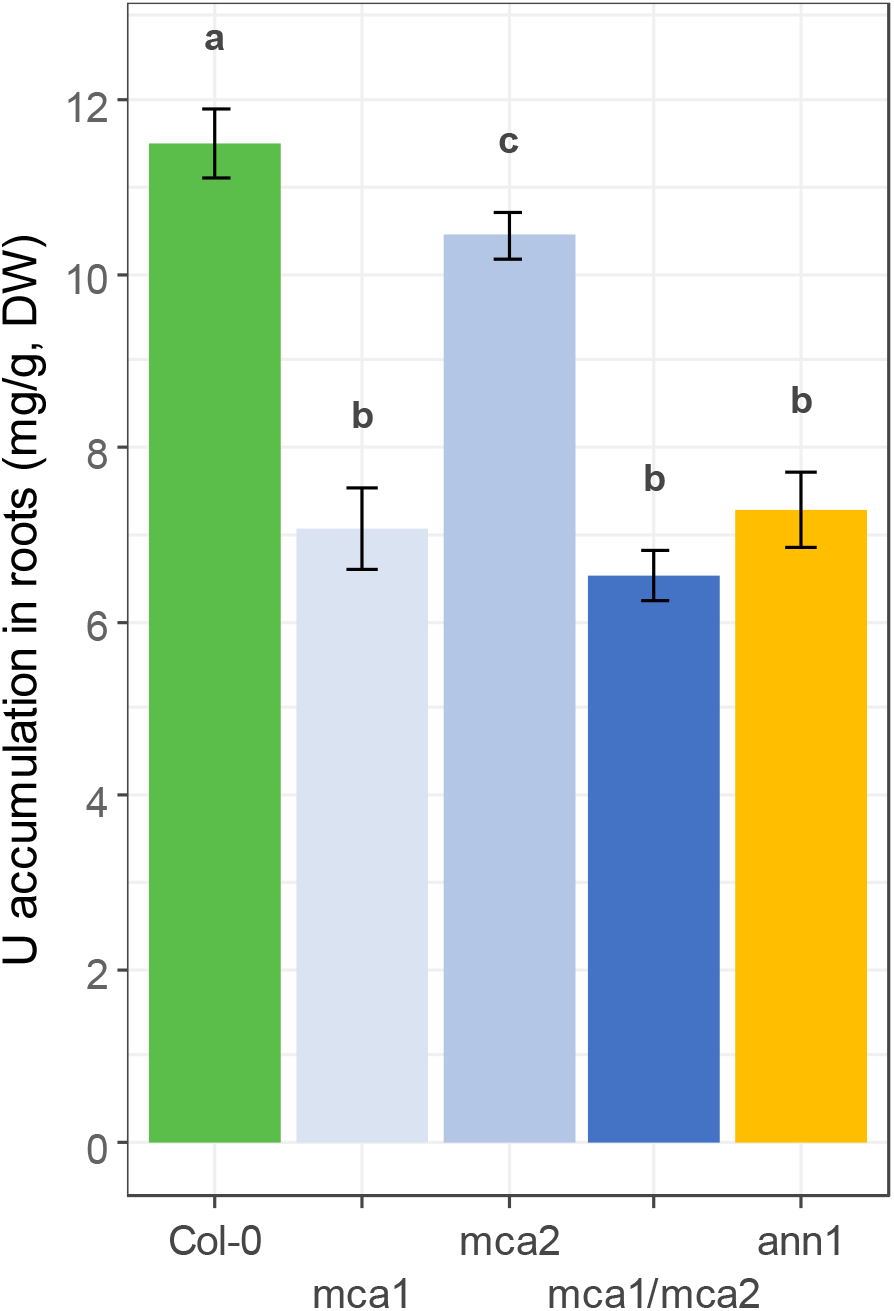
Bioaccumulation of U in the roots of Arabidopsis mutants deficient in Ca^2+^-permeable cation channels. The *A. thaliana* wild-type (Col-0) and the *mca1*, *mca2*, *mca1/mca2*, and *ann1* insertion mutants were grown for 4 weeks in Gre medium. Plants were challenged with 20 μM uranyl nitrate for 6 h and U (mg.g^−1^ dry weight) was measured by ICPMS. Bar plots represent mean ± SD with n=4 biological replicates. Letters indicate significant differences determined using a non-parametric Tukey’s test with p <0.01.

## Discussion

The link between U and the homeostasis of Ca has been previously reported in several plant species (Vanhoudt et al., 2011b; Tawussi et al., 2017; El Hayek et al., 2018; El Hayek et al., 2019; Rajabi et al., 2021) but no evidence was provided to support the uptake of the radionuclide by the Ca assimilation machinery. In this work, we provide several lines of evidence that U is taken up by the roots of Arabidopsis plants through Ca^2+^-permeable cation channels. The first evidence comes from U bioaccumulation assays with Arabidopsis plants starved for one or several essential nutrients. We postulated that changes in the expression of ion carriers located on the plasma membrane to adapt to nutrient starvation could modify U uptake and allow identification of the route(s) used by the radionuclide for absorption by roots. This assumption proved to be correct as our data showed that triple deprivation of Arabidopsis plants with Ca, K and Mg was accompanied with a 3-fold increase in the capacity of roots to accumulate U (Figure 1). Among the three cations, Ca was the only one for which starvation was correlated with an increased U accumulation. However, single Ca deprivation or double Ca-Mg or Ca-K deprivations were not as efficient as the triple Ca-Mg-K starvation to trigger U bioaccumulation (1.5-fold as compared with 3-fold increase; Figure 1). At least two hypotheses can explain this discrepancy, the magnitude of U accumulation in roots being related to i) the degree of Ca deficiency and/or ii) to a complex interplay between the homeostasis of Ca, Mg, and K. Under single and double Ca-deprivation conditions, Ca deficiency was much stronger than under triple Ca-Mg-K deprivation (30-60% and 20% reduction of Ca content in roots, respectively, and a comparable 40-50% decrease of Ca content in shoots; Figures S1 and S2, Tables S1 and S2). This suggests that the Ca influx machinery could not adapt to this drastic situation and, therefore, that U bioaccumulation could not be increased more than 1.5-fold. Also, crosstalk between nutrient homeostatic controls and the role of cytosolic free Ca in signaling several nutrient deficiencies, including K and Mg deficiency, are well documented (Tang et al., 2015; Wilkins et al., 2016; Tang and Luan, 2017; Bouain et al., 2019). Our ionomic analyses confirm the complex interaction between ions when nutrient supply is unbalanced (Figures S1 and S2, Tables S1 and S2) and suggest that the mechanisms implemented by Arabidopsis plants to adapt to the triple Ca-Mg-K deprivation, as yet unknown, lead to an optimal situation for the accumulation of U in roots.

The second line of evidence for the involvement of the Ca assimilation pathway in U uptake by roots comes from competition and inhibition assays. We showed that exogenous Ca inhibits U accumulation in roots in a dose-dependent manner (Figure 3), without modifying U bioavailability in the medium, suggesting a common route for the uptake of both cations. In addition, U and Ca have similar atomic radii (empirical values of 175 and 180 pm, respectively), supporting the view that they can compete for passing through the pore of Ca^2+^-permeable cation channels. This assumption is further supported by previous reports showing that U can interact strongly with Ca-binding proteins (e.g. calmodulin and osteopontin) (Qi et al., 2014; Brulfert et al., 2016; Creff et al., 2019), thanks to similar coordination properties of uranyl and Ca^2+^. Lanthanides (e.g. Gd^3+^), dihydropyridines (e.g. nifedipine) and phenylalkylamines (e.g. verapamil) are the most widely used blockers of Ca conductances in plants (De Vriese et al., 2018). Gadolinium was found to inhibit the bioaccumulation of U in Arabidopsis roots (Figure 4A) and tobacco BY2 cells (Rajabi et al., 2021), suggesting that MSCCs and NSCCs Ca^2+^-permeable channels serve as a possible route for U absorption in plant cells. Uranium bioaccumulation in Arabidopsis roots was also inhibited by nifedipine and verapamil (Figure 4B), two antagonists of VDCCs that bind to different receptor sites (De Vriese et al., 2018). The differences in time-dependent inhibition of U uptake by Gd^3+^ on the one hand and nifedipine and verapamil one the other hand (Figure 4) is likely due to the different modes of actions of the blockers (physical blocking of the channel pore by Gd^3+^, binding and inhibition for the drugs). Results obtained with Ca channels blockers should be interpreted with caution because side effects have been described (De Vriese et al., 2018). However, in combination with the other results described in this study, the pharmacological approach is in agreement with the involvement of different types of Ca^2+^-permeable cation channels (non-specific, mechanosensitive, voltage-dependent) in the uptake of U by root cells.

The structural, metabolic and signaling functions of Ca in plant cells are enabled by its orchestrated transport across cell membranes (Demidchik, 2018). Calcium influx across plant plasma membranes is mediated by highly redundant systems, including glutamate-like receptors (GLR, 20 genes in *A. thaliana*), cyclic nucleotide-gated channels (CNGC, 20 genes), mechanosensitive Ca^2+^-selective channels (MSCC, 10 genes), *mid1*-complementing activity channels (MCA, 2 genes), and the unconventional Ca channels annexins (8 genes) (Demidchik, 2018). In this study, we analyzed U bioaccumulation in null mutants for *MCA1*, *MCA2* and *ANNEXIN1* (*ANN1*) genes (Lee et al., 2004; Nakagawa et al., 2007; Yamanaka et al., 2010). We found that U bioaccumulation was reduced importantly (−40%) in *mca1*, *mca1*/*mca2* and *ann1*-null mutants, suggesting that these channels have a role in U uptake by roots (Figure 5). The MCA2 channel seems to have a limited role in U absorption (10% decrease as compared to the WT, Figure 5). MCA proteins from Arabidopsis can complement the Ca uptake deficiency in yeast cells lacking the high affinity Ca channel composed of the Mid1 and Cch1 subunits. The proteins are located at the plasma membrane and are inhibited by Gd^3+^ but not by verapamil (Nakagawa et al., 2007; Yamanaka et al., 2010). Although the role of MCA1 and MCA2 in Ca nutrition is still not clear, the expression patterns of the two genes suggest some distinct and overlapping functions. *MCA1* is expressed in the meristem and adjacent elongation zone of the primary root, detected in the stele and endodermis, but not in the cortex, epidermis, root cap, or root hairs. In the root, high levels of *MCA2* expression was found in vascular tissues, stele and endodermis, but no in the cortex, epidermis, root cap, meristem, elongation zone, or root hairs (Yamanaka et al., 2010). These patterns, together with the ability of the *mca1*, *mca2*, and *mca1/mca2* mutants to accumulate different amounts of U (Figure 5), suggest that MCA1 could contribute significantly to the symplastic transport of U in the elongation zone of the primary root whereas MCA1 and MCA2 could contribute to the radial transfer of U through the endodermis to reach vascular tissues. In our experimental conditions (acidic pH, absence of U chelators, 24 h of treatment), the translocation factor of U from roots to shoots is very low (2-7×10^−4^), meaning that the transfer of the radionuclide to the vascular tissues is poorly efficient and that a defect of MCA1 and MCA2 in the endodermis and stele has little impact on U bioaccumulation in roots. ANN1 is the predominantly abundant annexin in Arabidopsis and an *ann1*-null mutant displays phenotypes consistent with impaired Ca uptake (lack of Ca^2+^ conductance) at the root hair apex and epidermal elongation zone (Laohavisit et al., 2012). Our data suggest that ANN1 could be involved in the uptake of U in root hairs and the epidermal cell layer of the root elongation zone. Thus, MCA1 and ANN1 could contribute to the absorption of U in different zones and cell layers of the root. Additional Ca^2+^-permeable cation channels belonging to the NSCCs, MSCCs, and VDCCs are likely able to take part of U uptake in roots. To support this hypothesis and the postulate of our approach (nutrient depletion inducing the upregulation of ion carriers and, then, an improved U uptake), previous transcriptomic analyses have shown that several genes coding Ca^2+^-permeable channels are upregulated in Arabidopsis plants starved with Ca, of which glutamate-like receptors (*GLR1.2,1.4,2.5,2.8*), annexins (*ANN1,2,4,7*), mechanosensitive ion channel proteins (*MSL9,10*) and cyclic nucleotide-gated ion channels (*CNGC 9,11,19*) (Table S4) (Wang et al., 2015).

The partial inhibition (30 to 40%) of U accumulation in roots measured in competition and inhibition assays (Figures 3 and 4) or in mutants devoid of Ca^2+^-permeable cation channels (Figure 5) suggests that other nutrient assimilation pathways are involved and/or that U does not accumulate only inside root cells. Uranium bioaccumulation assays conducted at 4°C allowed us to discriminate between the fraction of the radionuclide associated with the symplast and the apoplast (Figure 2). The accumulation of U in the apoplast, which is a temperature-insensitive adsorption and diffusion process (Barberon and Geldner, 2014), accounts for 60-70% bioaccumulated U. This is in agreement with previous experiments showing that plant roots or plant cells exposed to U accumulate the radionuclide principally in the cell wall and intracellular spaces (Laurette et al., 2012a; Laurette et al., 2012b; El Hayek et al., 2018; El Hayek et al., 2019; Lai et al., 2020; Sarthou et al., 2020; Wu et al., 2020). The absorption of U by the symplastic pathway, which is mainly temperature-sensitive (reduction of energy metabolism, enzymatic activities, membrane fluidity, plasmodesmatal permeability) (Barberon and Geldner, 2014; Sager and Lee, 2014), accounts for 30-40% bioaccumulated U, and is virtually fully inhibited by external Ca at high concentration (Figure 3) or by inhibitors of NSCCs and Ca^2+^-permeable cation channels (Figure 4).

Our analyses of U bioaccumulation in nutrient-deprived plants provided no indication for the involvement of transport systems different from NSCCs or Ca^2+^-permeable cation channels in the absorption of U. In particular, Fe deficiency that is induced in the mi/100 medium (Figure S1, Table S2) and known to upregulate the expression of the IRT1 and NRAMP1 Fe^2+^ transporters (Curie et al., 2000; Vert et al., 2002; Castaings et al., 2016), was not accompanied by an improved bioaccumulation of U in roots. This finding confirms that, in an acidic environment devoid of U chelators, Fe^2+^ transporters are not involved in the uptake of U cations by roots (Berthet et al., 2018). Besides the Ca assimilation pathway, it is possible that endocytosis contributes to some extent to U uptake by root cells. Indeed, low-dose U uptake was found to be mainly mediated by endocytosis in kidney LLC-PK1 cells (Muller et al., 2008) and the pinocytotic uptake of U (high dose) in the root cap of oat seedlings could be observed (Wheeler and Hanchey, 1971). Also, it was found that deprivation of Ca in Arabidopsis roots enhanced endocytosis (Zhang et al., 2018), suggesting that the improved absorption of U in root cells of Ca-deficient plants could be due in part to this route.

## Conclusion

Understanding the molecular mechanisms governing the uptake of U is important for predicting its absorption by plants in different environments and for phytoremediation applications (Gavrilescu et al., 2009). Our study describes for the first time the involvement of Ca^2+^-permeable cation channels in the cellular uptake of U. In conditions favoring the formation of cationic species of U (acidic pH, low phosphate concentration, absence of carbonate or organic acids), the Ca assimilation pathway drives the main flux of the radionuclide within root cells. It is likely that, in environments with different U speciation, other transport systems are implicated in the uptake of U. For example, in carbonated water at nearly neutral pH, uranyl carbonate anionic forms are predominant and possibly incorporated into cells through anionic channels. Interestingly, in these conditions, Ca inhibits the radial transfer of U through the apoplastic pathway but facilitates the symplastic route and results in an increased translocation of U to shoot organs (El Hayek et al., 2018; El Hayek et al., 2019). Our study paves the way for the identification of U transport machinery in environmental conditions with changing U bioavailability as well as in other organisms, in which the molecular basis of U uptake is not known.

## Abbreviations

ANN: annexin
MCA: *mid1*-complementing activity

## Supporting information

## Authorship contribution statement

Manon Sarthou: conceptualization, investigation, writing - original draft, review and editing; Fabienne Devime: investigation; Célia Baggio: investigation; Sylvie Figuet: investigation; Claude Alban: investigation, writing - review and editing; Jacques Bourguignon: funding acquisition, conceptualization, investigation, writing - review and editing; Stéphane Ravanel: funding acquisition, conceptualization, investigation, writing - original draft, review and editing.

## Acknowledgements

We gratefully acknowledge Prof. Hidetoshi Iida (Tokyo Gakugei University) and Prof. Kenji Miura (University of Tsukuba) for kindly provided us with the *mca1* and *mca2* single and double mutants. We warmly acknowledge Dr. Stéphane Mari (INRAE, Montpellier) for stimulating discussions.

## Funding

This work was funded by the University Grenoble Alpes (PhD fellowship to Manon Sarthou), the Toxicology program of the CEA, the Plant Biology and Breeding division from INRAE, and the Agence Nationale de la Recherche (ANR-17-CE34-0007, GreenU project; ANR-17-EURE-0003, CBH-EUR-GS).

## Supporting information

**Table S1:** Composition of culture media and U speciation.

**1 - Composition of Gre media used for nutrient-deprivation experiments**
| | $\mu\text{M}$ | Gre | Gre/100 | M/100 | mi/100 | K/100 | Ca/100 | Mg/100 | KCa/100 | CaMg/100 | MgK/100 | P/100 |
| --- | --- | --- | --- | --- | --- | --- | --- | --- | --- | --- | --- | --- |
| Potassium sulfate | $\text{K}_2\text{SO}_4$ | 880 | 8.8 | 8.8 | 880 | 8.8 | 880 | 880 | 8.8 | 880 | 8.8 | 880 |
| Calcium nitrate • 4H<sub>2</sub>O | $\text{Ca}(\text{NO}_3)_2 \cdot 4\text{H}_2\text{O}$ | 1000 | 10 | 10 | 1000 | 1000 | 10 | 1000 | 10 | 10 | 1000 | 1000 |
| Magnesium sulfate • 7H<sub>2</sub>O | $\text{MgSO}_4 \cdot 7\text{H}_2\text{O}$ | 1000 | 10 | 10 | 1000 | 1000 | 1000 | 10 | 1000 | 10 | 10 | 1000 |
| Potassium phosphate monobasic | $\text{KH}_2\text{PO}_4$ | 250 | 2.5 | 2.5 | 250 | 2.5 | 250 | 250 | 2.5 | 250 | 2.5 | 2.5 |
| Boric acid | $\text{H}_3\text{BO}_3$ | 10 | 0.1 | 10 | 0.1 | 10 | 10 | 10 | 10 | 10 | 10 | 10 |
| Cupric sulfate • 5H<sub>2</sub>O | $\text{CuSO}_4 \cdot 5\text{H}_2\text{O}$ | 0.1 | 0.001 | 0.1 | 0.001 | 0.1 | 0.1 | 0.1 | 0.1 | 0.1 | 0.1 | 0.1 |
| Manganese sulfate • 1H<sub>2</sub>O | $\text{MnSO}_4 \cdot \text{H}_2\text{O}$ | 2 | 0.02 | 2 | 0.02 | 2 | 2 | 2 | 2 | 2 | 2 | 2 |
| Molybdic acid (ammonium salt) • 4H<sub>2</sub>O | $(\text{NH}_4)_6\text{Mo}_7\text{O}_{24} \cdot 4\text{H}_2\text{O}$ | 0.01 | 0.0001 | 0.01 | 0.0001 | 0.01 | 0.01 | 0.01 | 0.01 | 0.01 | 0.01 | 0.01 |
| Zinc sulfate • 7H<sub>2</sub>O | $\text{ZnSO}_4 \cdot 7\text{H}_2\text{O}$ | 2 | 0.02 | 2 | 0.02 | 2 | 2 | 2 | 2 | 2 | 2 | 2 |
| Sodium chloride | $\text{NaCl}$ | 10 | 0.1 | 10 | 0.1 | 10 | 10 | 10 | 10 | 10 | 10 | 10 |
| Cobalt chloride • 6H<sub>2</sub>O | $\text{CoCl}_2 \cdot 6\text{H}_2\text{O}$ | 0.02 | 0.0002 | 0.02 | 0.0002 | 0.02 | 0.02 | 0.02 | 0.02 | 0.02 | 0.02 | 0.02 |
| FeNaEDTA |  | 20 | 0.2 | 20 | 0.2 | 20 | 20 | 20 | 20 | 20 | 20 | 20 |
| MES |  | 250 | 250 | 250 | 250 | 250 | 250 | 250 | 250 | 250 | 250 | 250 |
| pH |  | 5.6 | 5.6 | 5.6 | 5.6 | 5.6 | 5.6 | 5.6 | 5.6 | 5.6 | 5.6 | 5.6 |
| Ammonium | $\text{NH}_4^+$ | 0.06 | 0.00 | 0.06 | 0.00 | 0.06 | 0.06 | 0.06 | 0.06 | 0.06 | 0.06 | 0.06 |
| Boric acid | $\text{H}_3\text{BO}_3$ | 10.00 | 0.10 | 10.00 | 0.10 | 10.00 | 10.00 | 10.00 | 10.00 | 10.00 | 10.00 | 10.00 |
| Calcium | $\text{Ca}^{2+}$ | 1 000.00 | 10.00 | 10.00 | 1 000.00 | 1 000.00 | 10.00 | 1 000.00 | 10.00 | 10.00 | 1 000.00 | 1 000.00 |
| Chloride | $\text{Cl}^-$ | 20.04 | 0.20 | 20.04 | 0.20 | 20.04 | 20.04 | 20.04 | 20.04 | 20.04 | 20.04 | 20.04 |
| Cobalt | $\text{Co}^{2+}$ | 0.02 | 0.00 | 0.02 | 0.00 | 0.02 | 0.02 | 0.02 | 0.02 | 0.02 | 0.02 | 0.02 |
| Copper | $\text{Cu}^{2+}$ | 0.10 | 0.00 | 0.10 | 0.00 | 0.10 | 0.10 | 0.10 | 0.10 | 0.10 | 0.10 | 0.10 |
| EDTA | EDTA | 20.00 | 0.20 | 20.00 | 0.20 | 20.00 | 20.00 | 20.00 | 20.00 | 20.00 | 20.00 | 20.00 |
| Fe II | $\text{Fe}^{2+}$ | 20.00 | 0.20 | 20.00 | 0.20 | 20.00 | 20.00 | 20.00 | 20.00 | 20.00 | 20.00 | 20.00 |
| Magnesium | $\text{Mg}^{2+}$ | 1 000.00 | 10.00 | 10.00 | 1 000.00 | 1 000.00 | 1 000.00 | 10.00 | 1 000.00 | 10.00 | 10.00 | 1 000.00 |
| Manganese | $\text{Mn}^{2+}$ | 2.00 | 0.02 | 2.00 | 0.02 | 2.00 | 2.00 | 2.00 | 2.00 | 2.00 | 2.00 | 2.00 |
| Molybdène | $\text{Mo}^{2+}$ | 0.07 | 0.00 | 0.07 | 0.00 | 0.07 | 0.07 | 0.07 | 0.07 | 0.07 | 0.07 | 0.07 |
| Nitrate | $\text{NO}_3^-$ | 2 000.00 | 20.00 | 20.00 | 2 000.00 | 2 000.00 | 20.00 | 2 000.00 | 20.00 | 20.00 | 2 000.00 | 2 000.00 |
| Phosphate | $\text{PO}_4^{3-}$ | 250.00 | 2.50 | 2.50 | 250.00 | 2.50 | 250.00 | 250.00 | 2.50 | 250.00 | 2.50 | 2.50 |
| Potassium | $\text{K}^+$ | 2 010.00 | 20.10 | 20.10 | 2 010.00 | 20.10 | 2 010.00 | 2 010.00 | 20.10 | 2 010.00 | 20.10 | 1 762.50 |
| Sodium | $\text{Na}^+$ | 10.00 | 0.10 | 10.00 | 0.10 | 10.00 | 10.00 | 10.00 | 10.00 | 10.00 | 10.00 | 10.00 |
| Sulfate | $\text{SO}_4^{2-}$ | 1 884.10 | 18.84 | 22.90 | 1 880.04 | 1 012.90 | 1 884.10 | 894.10 | 1 012.90 | 894.10 | 22.90 | 1 884.10 |
| Zinc | $\text{Zn}^{2+}$ | 2.00 | 0.02 | 2.00 | 0.02 | 2.00 | 2.00 | 2.00 | 2.00 | 2.00 | 2.00 | 2.00 |

**Table S1: Composition of culture media and U speciation****2 - Speciation of U in Gre media used for nutrient-deprivation experiments**Speciation was predicted using Visual MINTEQ (version 3.1, https://vminteq.lwr.kth.se/)**Concentration of U species (M)**
| Species name | Gre | Gre/100 | mi/100 | M/100 | Ca/100 | K/100 | Mg/100 | KCa/100 | MgK/100 | CaMg/100 | M/100 + (NH<sub>4</sub>)<sub>2</sub>SO<sub>4</sub> |
| --- | --- | --- | --- | --- | --- | --- | --- | --- | --- | --- | --- |
| (UO<sub>2</sub>)<sub>2</sub>(EDTA)2-4 | 1.47E-13 | 7.33E-16 | 1.43E-18 | 5.19E-14 | 1.75E-13 | 1.14E-13 | 1.04E-13 | 1.26E-13 | 7.10E-14 | 1.11E-13 | 1.19E-13 |
| (UO<sub>2</sub>)<sub>2</sub>(OH)<sub>2</sub>+2 | 6.40E-07 | 5.72E-07 | 6.85E-07 | 5.43E-07 | 6.01E-07 | 6.42E-07 | 6.30E-07 | 5.97E-07 | 6.26E-07 | 5.81E-07 | 5.83E-07 |
| (UO<sub>2</sub>)<sub>2</sub>EDTA (aq) | 1.85E-07 | 2.86E-08 | 5.99E-10 | 2.26E-07 | 2.25E-07 | 1.84E-07 | 1.86E-07 | 2.24E-07 | 1.82E-07 | 2.27E-07 | 2.27E-07 |
| (UO<sub>2</sub>)<sub>2</sub>OH+3 | 2.54E-09 | 1.49E-09 | 2.72E-09 | 1.45E-09 | 2.23E-09 | 2.40E-09 | 2.28E-09 | 2.04E-09 | 2.08E-09 | 1.90E-09 | 1.94E-09 |
| (UO<sub>2</sub>)<sub>2</sub>OHEDTA- | 5.42E-07 | 7.67E-08 | 1.75E-09 | 6.08E-07 | 6.47E-07 | 5.31E-07 | 5.32E-07 | 6.34E-07 | 5.12E-07 | 6.38E-07 | 6.40E-07 |
| (UO<sub>2</sub>)<sub>3</sub>(OH)<sub>4</sub>+2 | 4.70E-08 | 4.69E-08 | 5.20E-08 | 4.31E-08 | 4.39E-08 | 4.83E-08 | 4.77E-08 | 4.49E-08 | 4.88E-08 | 4.40E-08 | 4.39E-08 |
| (UO<sub>2</sub>)<sub>3</sub>(OH)<sub>5</sub>+ | 2.87E-06 | 3.68E-06 | 3.18E-06 | 3.34E-06 | 2.79E-06 | 3.06E-06 | 3.08E-06 | 3.00E-06 | 3.32E-06 | 3.02E-06 | 2.98E-06 |
| (UO<sub>2</sub>)<sub>3</sub>(OH)<sub>7</sub>- | 4.08E-12 | 5.24E-12 | 4.52E-12 | 4.76E-12 | 3.97E-12 | 4.35E-12 | 4.38E-12 | 4.27E-12 | 4.72E-12 | 4.29E-12 | 4.24E-12 |
| (UO<sub>2</sub>)<sub>4</sub>(OH)<sub>7</sub>+ | 4.82E-07 | 6.91E-07 | 5.53E-07 | 6.07E-07 | 4.67E-07 | 5.27E-07 | 5.32E-07 | 5.17E-07 | 5.92E-07 | 5.23E-07 | 5.14E-07 |
| UO<sub>2</sub>(OH)<sub>2</sub> (aq) | 6.34E-08 | 7.10E-08 | 6.56E-08 | 6.87E-08 | 6.32E-08 | 6.50E-08 | 6.54E-08 | 6.51E-08 | 6.74E-08 | 6.54E-08 | 6.51E-08 |
| UO<sub>2</sub>(OH)<sub>3</sub>- | 1.72E-10 | 1.76E-10 | 1.78E-10 | 1.71E-10 | 1.69E-10 | 1.74E-10 | 1.74E-10 | 1.71E-10 | 1.76E-10 | 1.70E-10 | 1.70E-10 |
| UO<sub>2</sub>(OH)<sub>4</sub>-2 | 6.11E-17 | 4.89E-17 | 6.32E-17 | 4.81E-17 | 5.77E-17 | 5.98E-17 | 5.85E-17 | 5.56E-17 | 5.64E-17 | 5.39E-17 | 5.44E-17 |
| UO<sub>2</sub>(SO<sub>4</sub>)<sub>2</sub>-2 | 2.34E-08 | 4.91E-12 | 2.40E-08 | 6.91E-12 | 2.82E-08 | 7.04E-09 | 6.37E-09 | 8.84E-09 | 4.46E-12 | 8.41E-09 | 1.06E-08 |
| UO<sub>2</sub>+2 | 1.79E-06 | 1.43E-06 | 1.85E-06 | 1.41E-06 | 1.69E-06 | 1.75E-06 | 1.71E-06 | 1.63E-06 | 1.65E-06 | 1.58E-06 | 1.59E-06 |
| UO<sub>2</sub>Cl+ | 1.70E-11 | 1.91E-13 | 1.76E-11 | 1.85E-11 | 1.70E-11 | 1.75E-11 | 1.76E-11 | 1.75E-11 | 1.81E-11 | 1.76E-11 | 1.75E-11 |
| UO<sub>2</sub>Cl<sub>2</sub> (aq) | 7.17E-18 | 9.57E-22 | 7.36E-18 | 9.19E-18 | 7.35E-18 | 7.53E-18 | 7.70E-18 | 7.84E-18 | 8.22E-18 | 8.05E-18 | 7.97E-18 |
| UO<sub>2</sub>EDTA-2 | 2.83E-08 | 2.79E-09 | 8.83E-11 | 2.31E-08 | 3.26E-08 | 2.61E-08 | 2.55E-08 | 2.95E-08 | 2.26E-08 | 2.87E-08 | 2.93E-08 |
| UO<sub>2</sub>H<sub>2</sub>PO<sub>4</sub>+ | 3.34E-09 | 3.60E-09 | 3.41E-09 | 3.52E-09 | 3.36E-09 | 3.39E-09 | 3.40E-09 | 3.42E-09 | 3.46E-09 | 3.43E-09 | 3.43E-09 |
| UO<sub>2</sub>H<sub>3</sub>PO<sub>4</sub>+2 | 4.67E-15 | 3.92E-15 | 4.77E-15 | 3.88E-15 | 4.51E-15 | 4.58E-15 | 4.51E-15 | 4.37E-15 | 4.35E-15 | 4.28E-15 | 4.32E-15 |
| UO<sub>2</sub>HEDTA- | 4.32E-08 | 5.47E-09 | 1.35E-10 | 4.49E-08 | 5.18E-08 | 4.13E-08 | 4.12E-08 | 4.93E-08 | 3.85E-08 | 4.93E-08 | 4.98E-08 |
| UO<sub>2</sub>HPO<sub>4</sub> (aq) | 6.26E-07 | 7.34E-07 | 6.39E-07 | 7.16E-07 | 6.38E-07 | 6.43E-07 | 6.50E-07 | 6.60E-07 | 6.73E-07 | 6.70E-07 | 6.67E-07 |
| UO<sub>2</sub>NO<sub>3</sub>+ | 5.37E-09 | 6.01E-11 | 5.55E-09 | 1.71E-10 | 1.57E-10 | 5.51E-09 | 5.52E-09 | 1.62E-10 | 5.70E-09 | 1.63E-10 | 1.62E-10 |
| UO<sub>2</sub>OH+ | 2.22E-06 | 2.29E-06 | 2.30E-06 | 2.22E-06 | 2.19E-06 | 2.25E-06 | 2.25E-06 | 2.21E-06 | 2.28E-06 | 2.20E-06 | 2.20E-06 |
| UO<sub>2</sub>PO<sub>4</sub>- | 1.27E-07 | 1.37E-07 | 1.30E-07 | 1.34E-07 | 1.28E-07 | 1.29E-07 | 1.30E-07 | 1.30E-07 | 1.32E-07 | 1.31E-07 | 1.31E-07 |
| UO<sub>2</sub>SO<sub>4</sub> (aq) | 1.65E-06 | 2.99E-08 | 1.70E-06 | 3.46E-08 | 1.86E-06 | 9.37E-07 | 9.05E-07 | 1.09E-06 | 2.52E-08 | 1.08E-06 | 1.21E-06 |

**Distribution of U species (%)**
| Species name | Gre | Gre/100 | mi/100 | M/100 | Ca/100 | K/100 | Mg/100 | KCa/100 | MgK/100 | CaMg/100 | M/100 + (NH<sub>4</sub>)<sub>2</sub>SO<sub>4</sub> |
| --- | --- | --- | --- | --- | --- | --- | --- | --- | --- | --- | --- |
| UO<sub>2</sub>+2 | 9.0 | 7.2 | 9.3 | 7.0 | 8.4 | 8.8 | 8.6 | 8.1 | 8.2 | 7.9 | 8.0 |
| UO<sub>2</sub>HEDTA- | 0.2 | 0.0 | - | 0.2 | 0.3 | 0.2 | 0.2 | 0.2 | 0.2 | 0.2 | 0.2 |
| UO<sub>2</sub>OH+ | 11.1 | 11.4 | 11.5 | 11.1 | 10.9 | 11.3 | 11.2 | 11.1 | 11.4 | 11.0 | 11.0 |
| (UO<sub>2</sub>)<sub>2</sub>(OH)<sub>2</sub>+2 | 6.4 | 5.7 | 6.8 | 5.4 | 6.0 | 6.4 | 6.3 | 6.0 | 6.3 | 5.8 | 5.8 |
| (UO<sub>2</sub>)<sub>3</sub>(OH)<sub>5</sub>+ | 43.0 | 55.2 | 47.7 | 50.2 | 41.9 | 45.8 | 46.1 | 45.0 | 49.7 | 45.3 | 44.7 |
| (UO<sub>2</sub>)<sub>4</sub>(OH)<sub>7</sub>+ | 9.6 | 13.8 | 11.1 | 12.2 | 9.3 | 10.5 | 10.6 | 10.3 | 11.8 | 10.5 | 10.3 |
| (UO<sub>2</sub>)<sub>3</sub>(OH)<sub>4</sub>+2 | 0.7 | 0.7 | 0.8 | 0.6 | 0.7 | 0.7 | 0.7 | 0.7 | 0.7 | 0.7 | 0.7 |
| (UO<sub>2</sub>)<sub>2</sub>OH+3 | 0.0 | 0.0 | 0.0 | 0.0 | 0.0 | 0.0 | 0.0 | 0.0 | 0.0 | 0.0 | 0.0 |
| UO<sub>2</sub>(OH)<sub>2</sub> (aq) | 0.3 | 0.4 | 0.3 | 0.3 | 0.3 | 0.3 | 0.3 | 0.3 | - | 0.3 | 0.3 |
| UO<sub>2</sub>SO<sub>4</sub> (aq) | 8.2 | 0.1 | 8.5 | 0.2 | 9.3 | 4.7 | 4.5 | 5.4 | 0.3 | 5.4 | 6.0 |
| UO<sub>2</sub>(SO<sub>4</sub>)<sub>2</sub>-2 | 0.1 | - | 0.1 | - | 0.1 | 0.0 | 0.0 | 0.0 | 0.1 | 0.0 | 0.1 |
| UO<sub>2</sub>NO<sub>3</sub>+ | 0.0 | - | 0.0 | - | - | 0.0 | 0.0 | - | 0.0 | - | - |
| UO<sub>2</sub>HPO<sub>4</sub> (aq) | 3.1 | 3.7 | 3.2 | 3.6 | 3.2 | 3.2 | 3.3 | 3.3 | 3.4 | 3.4 | 3.3 |
| UO<sub>2</sub>H<sub>2</sub>PO<sub>4</sub>+ | 0.0 | 0.0 | 0.0 | 0.0 | 0.0 | 0.0 | 0.0 | 0.0 | 0.0 | 0.0 | 0.0 |
| UO<sub>2</sub>PO<sub>4</sub>- | 0.6 | 0.7 | 0.6 | 0.7 | 0.6 | 0.6 | 0.6 | 0.7 | 0.7 | 0.7 | 0.7 |
| UO<sub>2</sub>EDTA-2 | 0.1 | 0.0 | - | 0.1 | 0.2 | 0.1 | 0.1 | 0.1 | 0.1 | 0.1 | 0.1 |
| (UO<sub>2</sub>)<sub>2</sub>OHEDTA- | 5.4 | 0.8 | 0.0 | 6.1 | 6.5 | 5.3 | 5.3 | 6.3 | 5.1 | 6.4 | 6.4 |
| (UO<sub>2</sub>)<sub>2</sub>EDTA (aq) | 1.9 | 0.3 | - | 2.3 | 2.2 | 1.8 | 1.9 | 2.2 | 1.8 | 2.3 | 2.3 |

Concentration of U species (M)
| Species name | Calcium nitrate (μM) |  |  |  |
| --- | --- | --- | --- | --- |
|  | 0 | 20 | 200 | 2000 |
| (UO<sub>2</sub>)<sub>2</sub>(OH)<sub>2</sub>+2 | 5.88E-07 | 5.95E-07 | 6.25E-07 | 7.28E-07 |
| (UO<sub>2</sub>)<sub>2</sub>OH+3 | 1.50E-09 | 1.55E-09 | 1.77E-09 | 2.69E-09 |
| (UO<sub>2</sub>)<sub>3</sub>(OH)<sub>4</sub>+2 | 4.94E-08 | 4.99E-08 | 5.19E-08 | 5.87E-08 |
| (UO<sub>2</sub>)<sub>3</sub>(OH)<sub>5</sub>+ | 3.93E-06 | 3.93E-06 | 3.89E-06 | 3.75E-06 |
| (UO<sub>2</sub>)<sub>3</sub>(OH)<sub>7</sub>- | 5.60E-12 | 5.59E-12 | 5.53E-12 | 5.33E-12 |
| (UO<sub>2</sub>)<sub>4</sub>(OH)<sub>7</sub>+ | 7.57E-07 | 7.53E-07 | 7.39E-07 | 6.92E-07 |
| UO<sub>2</sub>(OH)<sub>2</sub> (aq) | 7.27E-08 | 7.26E-08 | 7.19E-08 | 6.97E-08 |
| UO<sub>2</sub>(OH)<sub>3</sub>- | 1.80E-10 | 1.80E-10 | 1.82E-10 | 1.86E-10 |
| UO<sub>2</sub>(OH)<sub>4</sub>-2 | 4.91E-17 | 4.98E-17 | 5.28E-17 | 6.34E-17 |
| UO<sub>2</sub>+2 | 1.44E-06 | 1.46E-06 | 1.55E-06 | 1.86E-06 |
| UO<sub>2</sub>NO<sub>3</sub>+ | 1.21E-10 | 2.41E-10 | 1.31E-09 | 1.17E-08 |
| UO<sub>2</sub>OH+ | 2.33E-06 | 2.33E-06 | 2.35E-06 | 2.41E-06 |

Distribution of U species (%)
| Species name | Calcium nitrate (μM) |  |  |  |
| --- | --- | --- | --- | --- |
|  | 0 | 20 | 200 | 2000 |
| UO<sub>2</sub>+2 | 7.2 | 7.3 | 7.7 | 9.3 |
| UO<sub>2</sub>OH+ | 11.7 | 11.7 | 11.8 | 12.0 |
| (UO<sub>2</sub>)<sub>2</sub>(OH)<sub>2</sub>+2 | 5.9 | 5.9 | 6.2 | 7.3 |
| (UO<sub>2</sub>)<sub>3</sub>(OH)<sub>5</sub>+ | 59.0 | 58.9 | 58.3 | 56.2 |
| (UO<sub>2</sub>)<sub>4</sub>(OH)<sub>7</sub>+ | 15.1 | 15.1 | 14.8 | 13.8 |
| (UO<sub>2</sub>)<sub>3</sub>(OH)<sub>4</sub>+2 | 0.7 | 0.7 | 0.8 | 0.9 |
| (UO<sub>2</sub>)<sub>2</sub>OH+3 | 0.0 | 0.0 | 0.0 | 0.0 |
| UO<sub>2</sub>(OH)<sub>2</sub> (aq) | 0.4 | 0.4 | 0.4 | 0.3 |
| UO<sub>2</sub>NO<sub>3</sub>+ | - | - | - | 0.1 |

| **Concentration of U species (M)** |  |  | **Distribution of U species (%)** |  |  |
| --- | --- | --- | --- | --- | --- |
| **Species name** | **Control** | **GdCl<sub>3</sub> 250 µM** | **Species name** | **Control** | **GdCl<sub>3</sub> 250 µM** |
| (UO<sub>2</sub>)<sub>2</sub>(OH)<sub>2</sub>+2 | 5.88E-07 | 6.53E-07 | UO<sub>2</sub>+2 | 7.2 | 8.1 |
| (UO<sub>2</sub>)<sub>2</sub>OH+3 | 1.50E-09 | 1.99E-09 | UO<sub>2</sub>OH+ | 11.7 | 11.8 |
| (UO<sub>2</sub>)<sub>3</sub>(OH)<sub>4</sub>+2 | 4.94E-08 | 5.38E-08 | (UO<sub>2</sub>)<sub>2</sub>(OH)<sub>2</sub>+2 | 5.9 | 6.5 |
| (UO<sub>2</sub>)<sub>3</sub>(OH)<sub>5</sub>+ | 3.93E-06 | 3.85E-06 | (UO<sub>2</sub>)<sub>3</sub>(OH)<sub>5</sub>+ | 59.0 | 57.8 |
| (UO<sub>2</sub>)<sub>3</sub>(OH)<sub>7</sub>- | 5.60E-12 | 5.48E-12 | (UO<sub>2</sub>)<sub>4</sub>(OH)<sub>7</sub>+ | 15.1 | 14.5 |
| (UO<sub>2</sub>)<sub>4</sub>(OH)<sub>7</sub>+ | 7.57E-07 | 7.27E-07 | (UO<sub>2</sub>)<sub>3</sub>(OH)<sub>4</sub>+2 | 0.7 | 0.8 |
| UO<sub>2</sub>(OH)<sub>2</sub> (aq) | 7.27E-08 | 7.13E-08 | (UO<sub>2</sub>)<sub>2</sub>OH+3 | 0.0 | 0.0 |
| UO<sub>2</sub>(OH)<sub>3</sub>- | 1.80E-10 | 1.83E-10 | UO<sub>2</sub>(OH)<sub>2</sub> (aq) | 0.4 | 0.4 |
| UO<sub>2</sub>(OH)<sub>4</sub>-2 | 4.91E-17 | 5.56E-17 |  |  |  |
| UO<sub>2</sub>+2 | 1.44E-06 | 1.63E-06 |  |  |  |
| UO<sub>2</sub>Cl+ | - | 1.43E-09 |  |  |  |
| UO<sub>2</sub>Cl<sub>2</sub> (aq) | - | 5.04E-14 |  |  |  |
| UO<sub>2</sub>NO<sub>3</sub>+ | 1.21E-10 | 1.18E-10 |  |  |  |
| UO<sub>2</sub>OH+ | 2.33E-06 | 2.37E-06 |  |  |  |

**Table S2:** Root ionomes after 4 days of nutrient deprivations. Four-week-old Arabidopsis plants grown in a complete standard Gre medium were transferred for 4 days in Gre medium depleted of one or several nutrients (100-fold reduction of the standard concentration; Table S1). Element concentrations (in μg.g^−1^ dry weight) were measured in mineralized roots by ICP-MS. Deprivation experiments have been performed from 2 to 11 times. The table is a representative root ionome from two independent experiments with n=4 plants. Letters indicate significant differences determined using a non-parametric Tukey’s test with p<0.01.

| Element<br>($\mu\text{g/g}$) | Culture medium | | | | | | | | | |
| --- | --- | --- | --- | --- | --- | --- | --- | --- | --- | --- |
|  | Gre | Gre/100 | M/100 | mi/100 | K/100 | Ca/100 | Mg/100 | KCa/100 | CaMg/100 | MgK/100 |
| **Mg** | 2960 ± 91<sup>a</sup> | 2300 ± 76<sup>b</sup> | 2230 ± 26<sup>b</sup> | 3017 ± 40<sup>a</sup> | 3583 ± 33<sup>c</sup> | 5864 ± 164<sup>d</sup> | 1897 ± 26<sup>e</sup> | 4804 ± 142<sup>f</sup> | 1769 ± 51<sup>e</sup> | 2447 ± 71<sup>b</sup> |
| **P** | 16170 ± 541<sup>a</sup> | 10920 ± 334<sup>b</sup> | 11623 ± 239<sup>b</sup> | 14287 ± 113<sup>c</sup> | 9181 ± 446<sup>e</sup> | 18167 ± 378<sup>d</sup> | 13139 ± 321<sup>f</sup> | 9645 ± 243<sup>e</sup> | 14938 ± 74<sup>a</sup> | 9590 ± 212<sup>e</sup> |
| **K** | 69658 ± 1910<sup>a,b</sup> | 51856 ± 1553<sup>e,c</sup> | 50002 ± 546<sup>e</sup> | 72040 ± 2147<sup>a</sup> | 64787 ± 2119<sup>b</sup> | 55029 ± 2160<sup>c</sup> | 73187 ± 1127<sup>a</sup> | 62262 ± 1611<sup>b</sup> | 50673 ± 822<sup>e,c</sup> | 46649 ± 1115<sup>d</sup> |
| **Ca** | 4634 ± 93<sup>a,b</sup> | 3590 ± 139<sup>e,c</sup> | 3690 ± 152<sup>e,c</sup> | 3865 ± 104<sup>e</sup> | 3916 ± 181<sup>a,e</sup> | 2126 ± 149<sup>d</sup> | 4459 ± 126<sup>a,b</sup> | 1721 ± 108<sup>f</sup> | 3150 ± 187<sup>c</sup> | 5334 ± 217<sup>b</sup> |
| **Mn** | 78 ± 9<sup>a,b</sup> | 32 ± 2<sup>e</sup> | 161 ± 8<sup>f</sup> | 32 ± 5<sup>e</sup> | 56 ± 10<sup>a,c</sup> | 88 ± 18<sup>a,b,d</sup> | 84 ± 4<sup>b</sup> | 43 ± 5<sup>c</sup> | 194 ± 8<sup>g</sup> | 121 ± 3<sup>d</sup> |
| **Fe** | 6854 ± 85<sup>a</sup> | 4325 ± 201<sup>b</sup> | 5533 ± 110<sup>e</sup> | 4223 ± 93<sup>b</sup> | 6120 ± 460<sup>a,e,c</sup> | 6901 ± 163<sup>a</sup> | 4696 ± 245<sup>b</sup> | 4136 ± 213<sup>b</sup> | 5390 ± 175<sup>e</sup> | 6282 ± 216<sup>c</sup> |
| **Zn** | 795 ± 32<sup>a,b</sup> | 597 ± 28<sup>e</sup> | 945 ± 40<sup>c</sup> | 638 ± 63<sup>e,d,f</sup> | 805 ± 24<sup>a,d</sup> | 474 ± 42<sup>f</sup> | 631 ± 29<sup>e</sup> | 680 ± 43<sup>b,e</sup> | 616 ± 50<sup>e</sup> | 1064 ± 84<sup>c</sup> |
| **Mo** | 75 ± 3<sup>a,b</sup> | 52 ± 4<sup>e</sup> | 65 ± 2<sup>c</sup> | 42 ± 3<sup>e</sup> | 96 ± 5<sup>d,f</sup> | 69 ± 5<sup>a,c</sup> | 78 ± 4<sup>a,b,c,d</sup> | 110 ± 5<sup>f</sup> | 37 ± 2<sup>e</sup> | 107 ± 8<sup>b,d,f</sup> |

**Table S3:** Shoot ionomes after 4 days of nutrient deprivations. Four-week-old Arabidopsis plants grown in a complete standard Gre medium were transferred for 4 days in Gre medium depleted of one or several nutrients (100-fold reduction of the standard concentration; Table S1). Element concentrations (in μg.g^−1^ dry weight) were measured in mineralized shoots by ICP-MS. Deprivation experiments have been performed from 2 to 11 times. The table is a representative shoot ionome from two independent experiments with n=4 plants. Letters indicate significant differences determined using a non-parametric Tukey’s test with p<0.01.

| Element<br>($\mu\text{g/g}$) | Culture medium | | | | | | | | | |
| --- | --- | --- | --- | --- | --- | --- | --- | --- | --- | --- |
|  | Gre | Gre/100 | M/100 | mi/100 | K/100 | Ca/100 | Mg/100 | KCa/100 | CaMg/100 | MgK/100 |
| **Mg** | 10687 ± 93<sup>a</sup> | 7702 ± 131<sup>b</sup> | 6880 ± 83<sup>c</sup> | 11078 ± 129<sup>d</sup> | 11010 ± 82<sup>a,d</sup> | 9440 ± 320<sup>e</sup> | 6561 ± 134<sup>f</sup> | 10672 ± 232<sup>a,d,e</sup> | 5467 ± 359<sup>f</sup> | 2891 ± 3<sup>g</sup> |
| **P** | 6606 ± 243<sup>a</sup> | 5026 ± 128<sup>b</sup> | 4170 ± 113<sup>c</sup> | 6482 ± 191<sup>a</sup> | 3948 ± 96<sup>c</sup> | 6307 ± 130<sup>a</sup> | 6041 ± 160<sup>a</sup> | 4763 ± 117<sup>b</sup> | 5725 ± 257<sup>a</sup> | 2061 ± 25<sup>d</sup> |
| **K** | 30506 ± 608<sup>a</sup> | 29067 ± 277<sup>b</sup> | 24720 ± 521<sup>c</sup> | 29044 ± 888<sup>a,b</sup> | 25269 ± 1026<sup>c</sup> | 25054 ± 821<sup>c,d</sup> | 31305 ± 500<sup>a</sup> | 22972 ± 360<sup>d</sup> | 27674 ± 1262<sup>a,b,c</sup> | 13947 ± 269<sup>e</sup> |
| **Ca** | 28023 ± 868<sup>a</sup> | 17499 ± 746<sup>b</sup> | 14008 ± 386<sup>c</sup> | 26339 ± 593<sup>a</sup> | 26374 ± 536<sup>a</sup> | 14786 ± 222<sup>c,d</sup> | 32479 ± 871<sup>e</sup> | 16695 ± 989<sup>b,d</sup> | 14386 ± 546<sup>c,d</sup> | 17292 ± 141<sup>b</sup> |
| **Mn** | 100 ± 3<sup>a</sup> | 72 ± 3<sup>b,c,d</sup> | 127 ± 2<sup>e</sup> | 65 ± 2<sup>b</sup> | 99 ± 2<sup>a</sup> | 82 ± 4<sup>c</sup> | 118 ± 3<sup>f</sup> | 98 ± 3<sup>a</sup> | 114 ± 3<sup>f</sup> | 69 ± 1<sup>d</sup> |
| **Fe** | 73 ± 2<sup>a</sup> | 75 ± 8<sup>a</sup> | 61 ± 2<sup>b</sup> | 62 ± 5<sup>a,b</sup> | 73 ± 1<sup>a</sup> | 52 ± 0<sup>c</sup> | 70 ± 3<sup>a</sup> | 72 ± 3<sup>a</sup> | 57 ± 16<sup>a,b</sup> | 37 ± 1<sup>d</sup> |
| **Zn** | 110 ± 5<sup>a,b</sup> | 124 ± 5<sup>a,b,c</sup> | 129 ± 8<sup>a,c</sup> | 89 ± 4<sup>d</sup> | 118 ± 4<sup>a</sup> | 104 ± 6<sup>b</sup> | 112 ± 2<sup>b</sup> | 149 ± 8<sup>c</sup> | 124 ± 7<sup>a,b</sup> | 74 ± 5<sup>d</sup> |
| **Mo** | 14,5 ± 0,4<sup>a</sup> | 11,7 ± 0,9<sup>b</sup> | 11,1 ± 0,5<sup>b</sup> | 10,7 ± 0,3<sup>b</sup> | 14,3 ± 0,7<sup>a,c</sup> | 11,5 ± 1,2<sup>b,d</sup> | 14,8 ± 0,5<sup>a</sup> | 12,1 ± 0,5<sup>b,c</sup> | 10,7 ± 0,5<sup>b</sup> | 7,5 ± 0,3<sup>d</sup> |

**Table S4:**
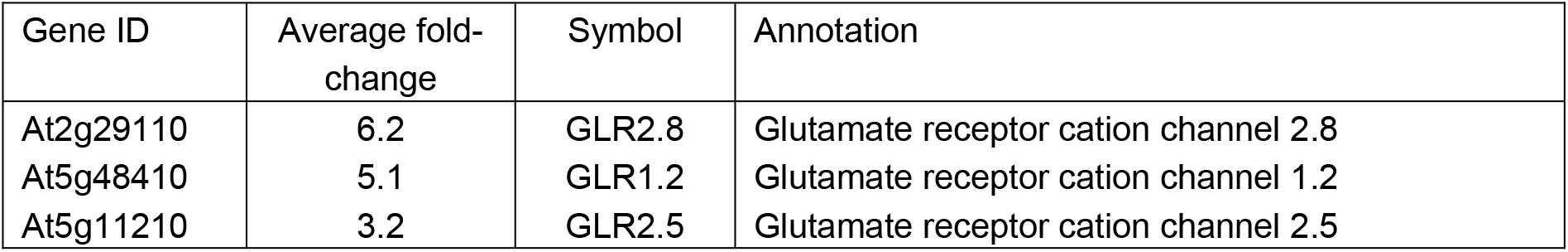
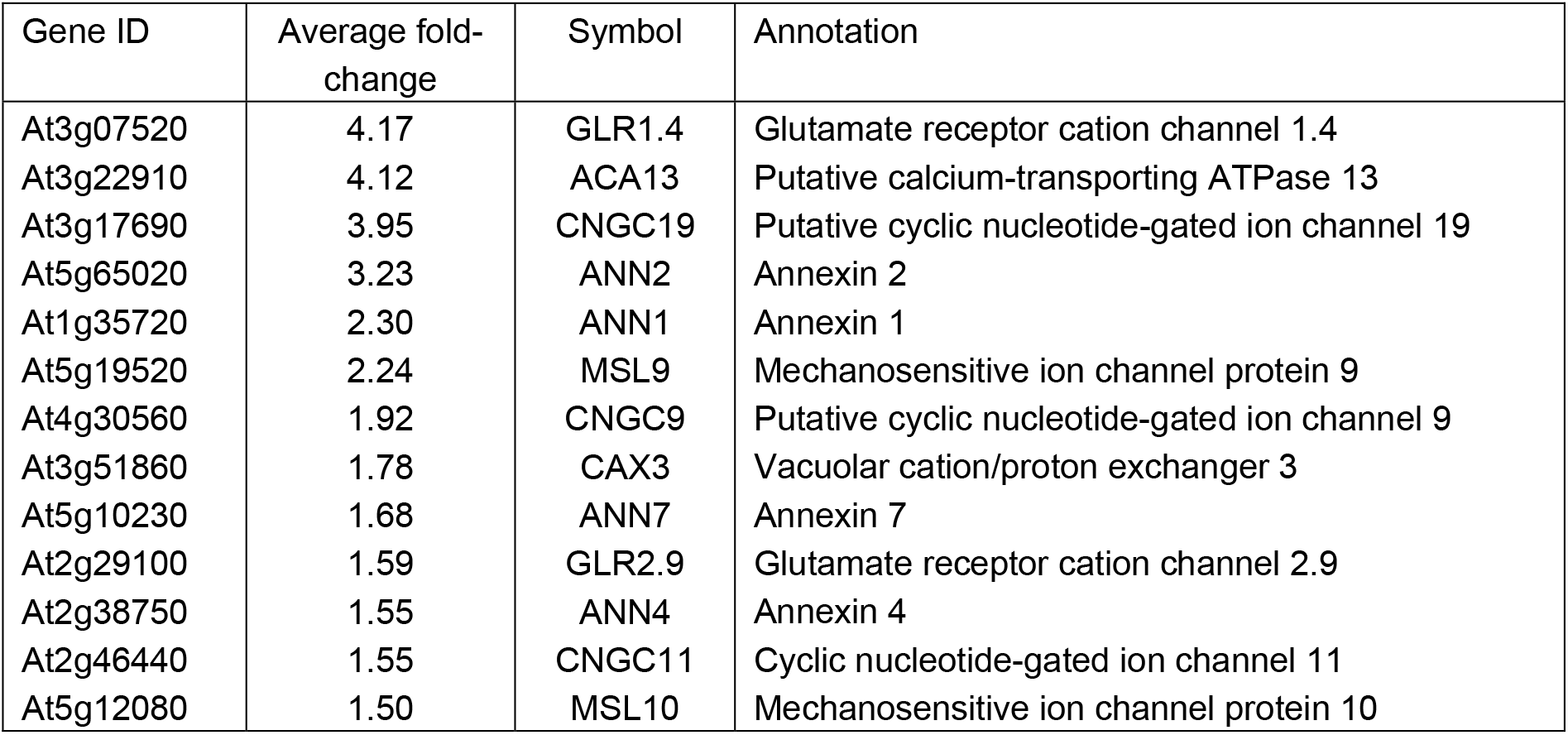
Up-regulation of genes coding calcium channels and transporters in response to extracellular calcium depletion. - <u>Data from Wang *et al* (2015)</u> Arabidopsis transcriptional response to extracellular Ca^2+^ depletion involves a transient rise in cytosolic Ca^2+^. *J Int Plant Biol 57*, 138-150. Upregulated genes involved in Ca mobilization (24 h Ca depletion in 11-day-old *Arabidopsis thaliana* seedlings). Data from Table 2.
- <u>Transcriptomic data GSM1655511 (unpublished).</u> *Arabidopsis thaliana* plants were grown on perlite with a complete nutrient solution in which Ca concentration was 3 mM. After 17 days Ca was suppressed for 2 days. Upregulated genes in Ca-deficient plants (>1.5-fold change)

| Gene ID | Average fold-change | Symbol | Annotation |
| --- | --- | --- | --- |
| At2g29110 | 6.2 | GLR2.8 | Glutamate receptor cation channel 2.8 |
| At5g48410 | 5.1 | GLR1.2 | Glutamate receptor cation channel 1.2 |
| At5g11210 | 3.2 | GLR2.5 | Glutamate receptor cation channel 2.5 |
| At3g07520 | 4.17 | GLR1.4 | Glutamate receptor cation channel 1.4 |
| At3g22910 | 4.12 | ACA13 | Putative calcium-transporting ATPase 13 |
| At3g17690 | 3.95 | CNGC19 | Putative cyclic nucleotide-gated ion channel 19 |
| At5g65020 | 3.23 | ANN2 | Annexin 2 |
| At1g35720 | 2.30 | ANN1 | Annexin 1 |
| At5g19520 | 2.24 | MSL9 | Mechanosensitive ion channel protein 9 |
| At4g30560 | 1.92 | CNGC9 | Putative cyclic nucleotide-gated ion channel 9 |
| At3g51860 | 1.78 | CAX3 | Vacuolar cation/proton exchanger 3 |
| At5g10230 | 1.68 | ANN7 | Annexin 7 |
| At2g29100 | 1.59 | GLR2.9 | Glutamate receptor cation channel 2.9 |
| At2g38750 | 1.55 | ANN4 | Annexin 4 |
| At2g46440 | 1.55 | CNGC11 | Cyclic nucleotide-gated ion channel 11 |
| At5g12080 | 1.50 | MSL10 | Mechanosensitive ion channel protein 10 |

**Figure S1:**
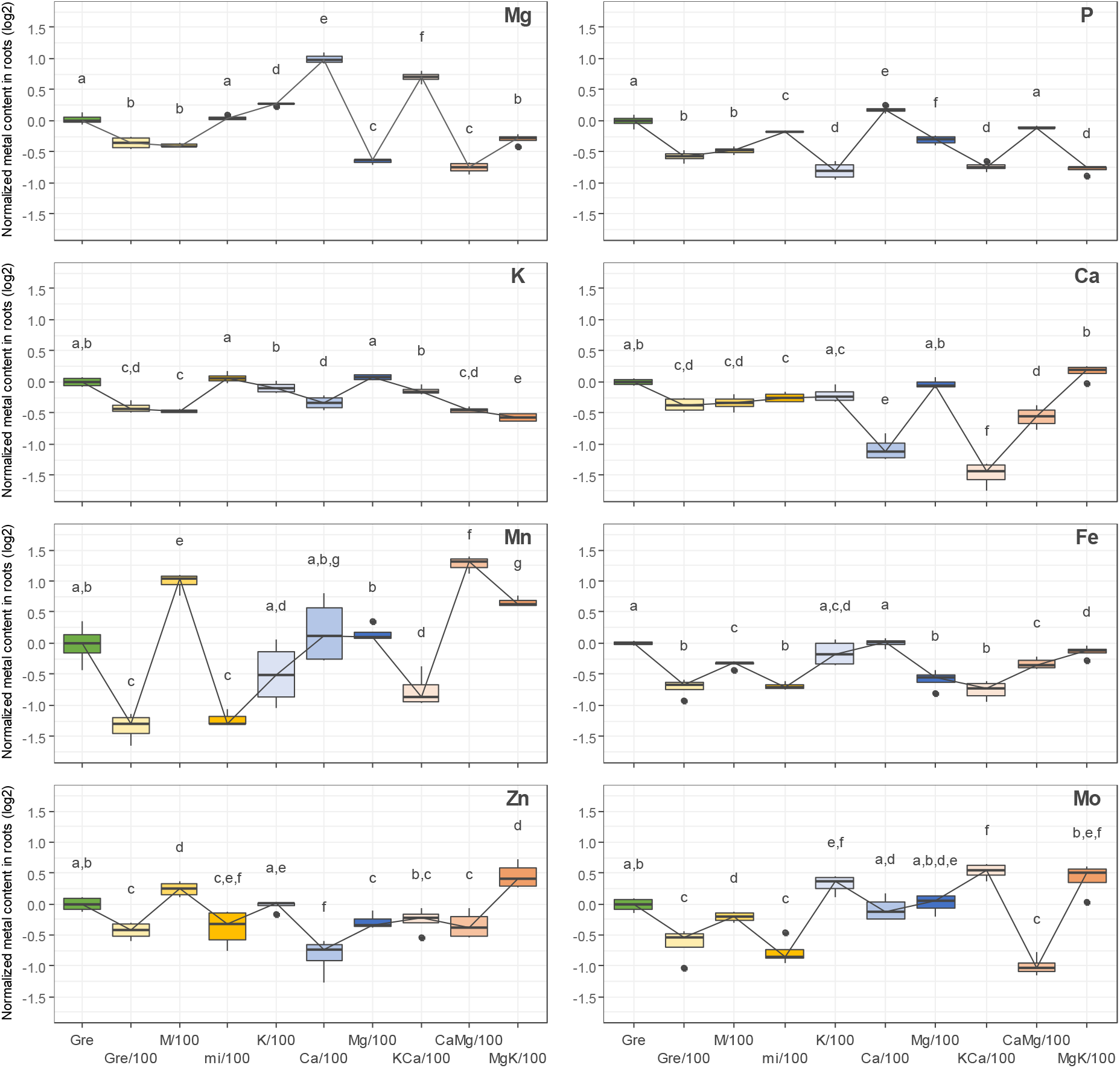
Changes in root ionomes induced by 4 days of nutrient deprivations. Modifications of the root ionomes from two independent representative deprivation experiments (Table S2) are shown. Changes in mineral content are expressed as a log2 of the ratio relative to the control condition (Gre medium). Box plots represent minimum, first quartile, median, third quartile and maximum. Letters indicate significant differences determined using a non-parametric Tukey’s test with p<0.01.

**Figure S2:**
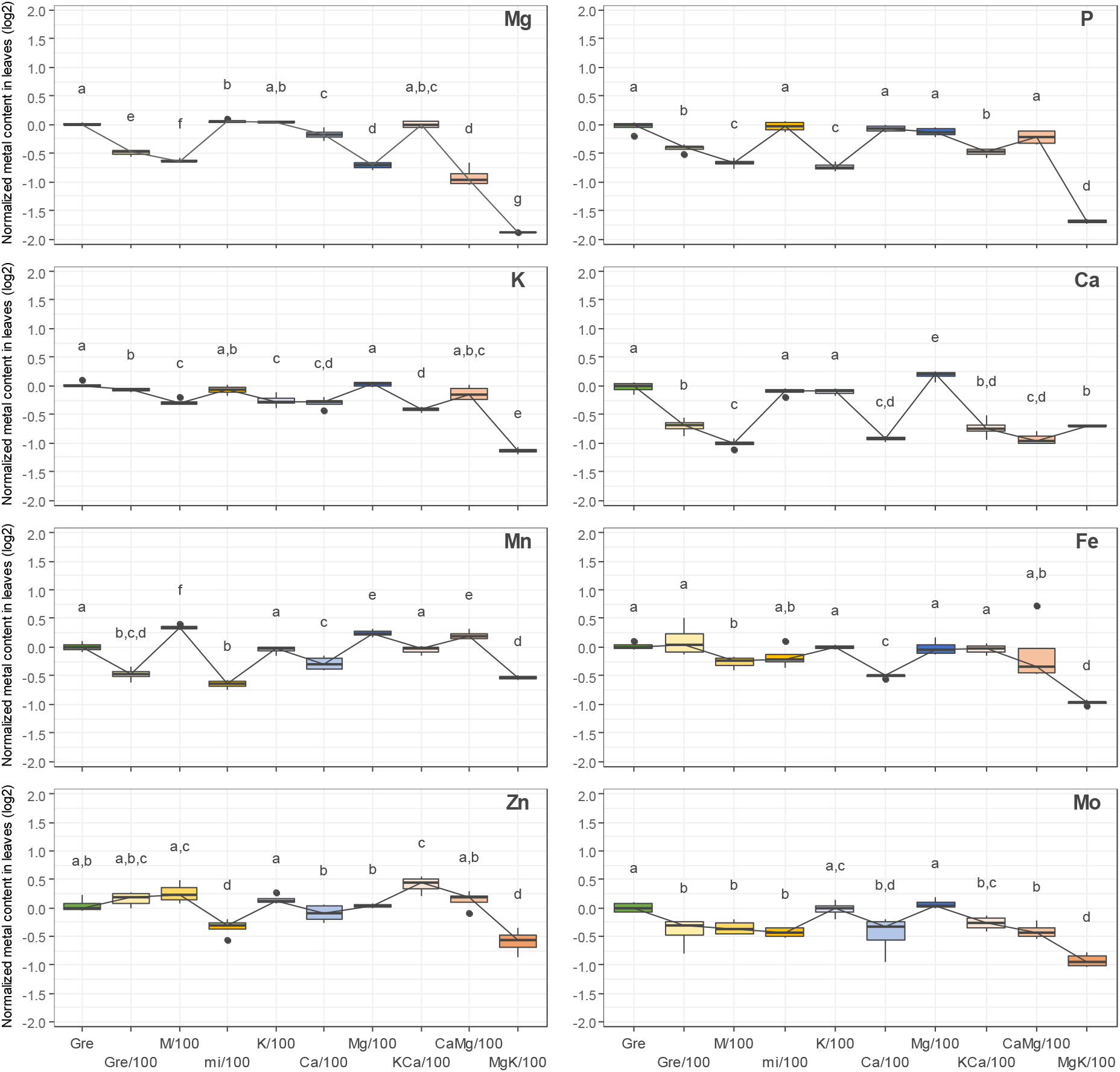
Changes in shoot ionomes induced by 4 days of nutrient deprivations. Modifications of the shoot ionomes from two independent representative deprivation experiments (Table S3) are shown. Changes in mineral content are expressed as a log2 of the ratio relative to the control condition (Gre medium). Box plots represent minimum, first quartile, median, third quartile and maximum. Letters indicate significant differences determined using a non-parametric Tukey’s test with p<0.01.

**Figure S3:**
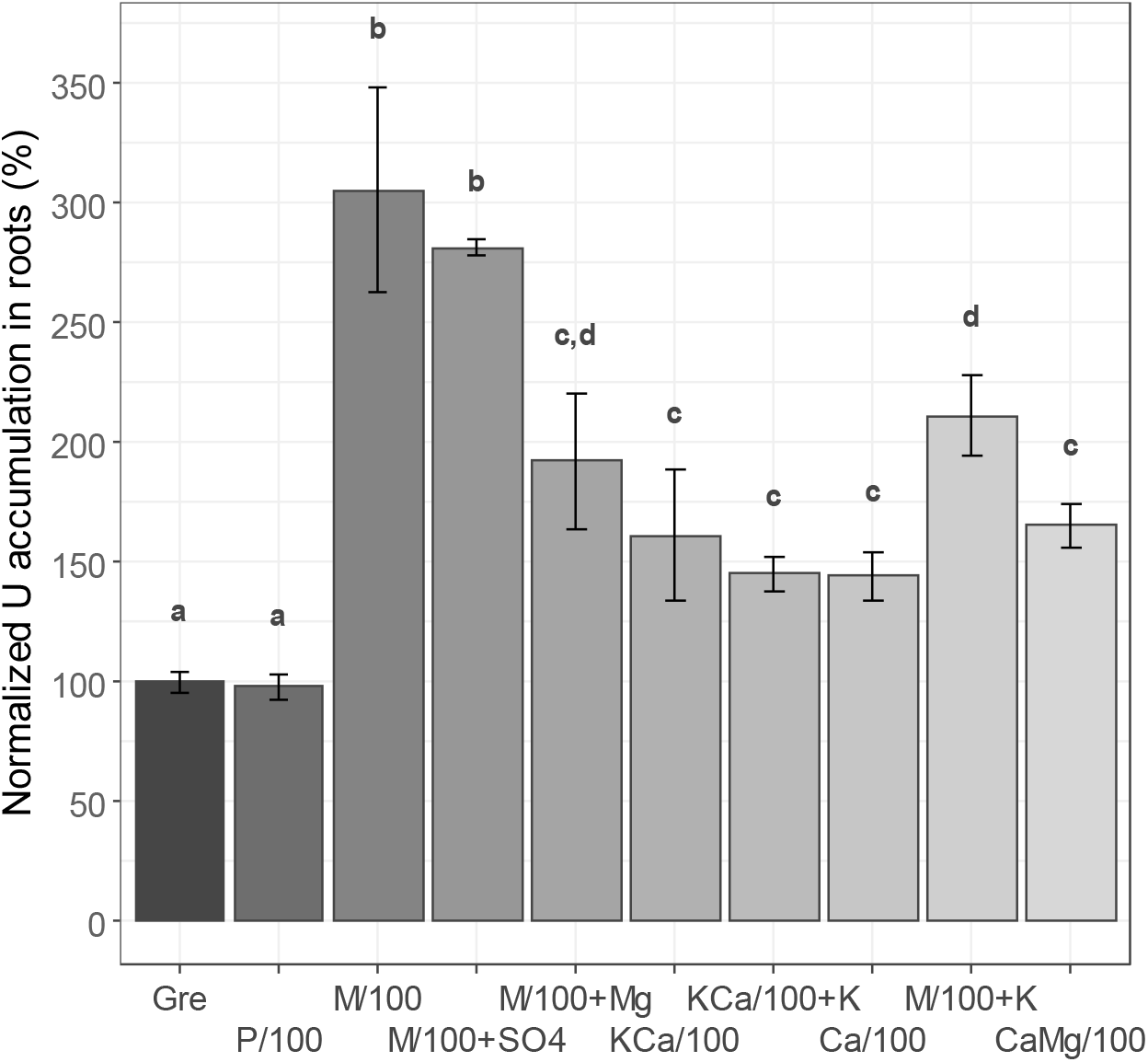
Bioaccumulation of U in the roots of Arabidopsis plants deprived with nutrients, effects of supplementation with anions. Four-week-old Arabidopsis plants grown in a complete standard Gre medium were transferred for 4 days in Gre medium depleted of one or several nutrients (100-fold reduction of the standard concentration; Table S1). Various media have been tested to modify the nutrient supply of plants. “P/100” is a Gre medium in which phosphate has been reduced by 100-fold. “M/100+SO4” is a M/100 medium supplemented with 1 mM (NH_4_)_2_SO_4_ to restore sulfate supply to control conditions. “KCa/100+K” is a KCa/100 medium supplemented with 2 mM KNO_3_, leading to a medium comparable to Ca/100 but with a standard nitrate concentration. “M/100+K” is the M/100 medium with 2 mM KNO_3_, leading to a medium comparable to CaMg/100 but with a standard nitrate concentration. Nutrient-deprived plants were challenged with 20 μM uranyl nitrate for 24 h, thoroughly washed with sodium carbonate and distilled water to remove loosely-bound metals, and U was measured by ICPMS in mineralized roots. Uranium accumulation in roots was expressed as a percentage related to the control condition in Gre medium. Bar plots represent mean ± SD. Letters indicate significant differences determined using a non-parametric Tukey’s test with p<0.01. The amount of U bioaccumulated in Gre medium was 727 ± 79 μg.g^−1^ fresh weight (mean ± SD, n=11 independent experiments).

## Notes

### Competing Interest Statement

The authors have declared no competing interest.

## References

Anke M, Seeber O, Müller R, Schäfer U, Zerull J (2009) Uranium transfer in the food chain from soil to plants, animals and man. Geochemistry 69: 75–90

Aranjuelo I, Doustaly F, Cela J, Porcel R, Müller M, Aroca R, Munné-Bosch S, Bourguignon J (2014) Glutathione and transpiration as key factors conditioning oxidative stress in Arabidopsis thaliana exposed to uranium. Planta 239: 817–830

Barberon M, Geldner N (2014) Radial Transport of Nutrients: The Plant Root as a Polarized Epithelium. Plant Physiology 166: 528–537

Berthet S, Villiers F, Alban C, Serre NBC, Martin-Laffon J, Figuet S, Boisson A-M, Bligny R, Kuntz M, Finazzi G, et al (2018) Arabidopsis thaliana plants challenged with uranium reveal new insights into iron and phosphate homeostasis. New Phytologist 217: 657–670

Bouain N, Krouk G, Lacombe B, Rouached H (2019) Getting to the Root of Plant Mineral Nutrition: Combinatorial Nutrient Stresses Reveal Emergent Properties. Trends in Plant Science 24: 542–552

Brulfert F, Safi S, Jeanson A, Martinez-Baez E, Roques J, Berthomieu C, Solari P-L, Sauge-Merle S, Simoni É (2016) Structural Environment and Stability of the Complexes Formed Between Calmodulin and Actinyl Ions. Inorg Chem 55: 2728–2736

Castaings L, Caquot A, Loubet S, Curie C (2016) The high-affinity metal Transporters NRAMP1 and IRT1 Team up to Take up Iron under Sufficient Metal Provision. Scientific Reports 6: 37222

Chen L, Liu J, Zhang W, Zhou J, Luo D, Li Z (2021) Uranium (U) source, speciation, uptake, toxicity and bioremediation strategies in soil-plant system: A review. Journal of Hazardous Materials 413: 125319

Creff G, Zurita C, Jeanson A, Carle G, Vidaud C, Auwer CD (2019) What do we know about actinides-proteins interactions? Radiochimica Acta 107: 993–1009

Curie C, Alonso JM, Jean ML, Ecker JR, Briat J-F (2000) Involvement of NRAMP1 from Arabidopsis thaliana in iron transport. Biochemical Journal 347: 749–755

Davies JM (2014) Annexin-Mediated Calcium Signalling in Plants. Plants (Basel) 3: 128–140

De Vriese K, Costa A, Beeckman T, Vanneste S (2018) Pharmacological Strategies for Manipulating Plant Ca2+ Signalling. International Journal of Molecular Sciences 19: 1506

Demidchik V (2018) ROS-Activated Ion Channels in Plants: Biophysical Characteristics, Physiological Functions and Molecular Nature. Int J Mol Sci 19: 1263

Doustaly F, Combes F, Fiévet JB, Berthet S, Hugouvieux V, Bastien O, Aranjuelo I, Leonhardt N, Rivasseau C, Carrière M, et al (2014) Uranium perturbs signaling and iron uptake response in Arabidopsis thaliana roots. Metallomics 6: 809–821

Ebbs SD, Brady DJ, Kochian LV (1998) Role of uranium speciation in the uptake and translocation of uranium by plants. Journal of Experimental Botany 49: 1183–1190

El Hayek E, Brearley AJ, Howard T, Hudson P, Torres C, Spilde MN, Cabaniss S, Ali A-MS, Cerrato JM (2019) Calcium in Carbonate Water Facilitates the Transport of U(VI) in Brassica juncea Roots and Enables Root-to-Shoot Translocation. ACS Earth Space Chem 3: 2190–2196

El Hayek E, Torres C, Rodriguez-Freire L, Blake JM, De Vore CL, Brearley AJ, Spilde MN, Cabaniss S, Ali A-MS, Cerrato JM (2018) Effect of Calcium on the Bioavailability of Dissolved Uranium(VI) in Plant Roots under Circumneutral pH. Environmental Science & Technology 52: 13089–13098

Fortin C, Denison FH, Garnier-Laplace J (2007) Metal-phytoplankton interactions: Modeling the effect of competing ions (H+, Ca2+, and Mg2+) on uranium uptake. Environmental Toxicology and Chemistry 26: 242–248

Gao N, Huang Z, Liu H, Hou J, Liu X (2019) Advances on the toxicity of uranium to different organisms. Chemosphere 237: 124548

Gavrilescu M, Pavel LV, Cretescu I (2009) Characterization and remediation of soils contaminated with uranium. Journal of Hazardous Materials 163: 475–510

Kiba T, Feria-Bourrellier A-B, Lafouge F, Lezhneva L, Boutet-Mercey S, Orsel M, Bréhaut V, Miller A, Daniel-Vedele F, Sakakibara H, et al (2012) The Arabidopsis Nitrate Transporter NRT2.4 Plays a Double Role in Roots and Shoots of Nitrogen-Starved Plants. The Plant Cell 24: 245–258

Konietschke F, Placzek M, Schaarschmidt F, Hothorn LA (2015) nparcomp: An R Software Package for Nonparametric Multiple Comparisons and Simultaneous Confidence Intervals. Journal of Statistical Software 64: 1–17

Lai J, Liu Z, Li C, Luo X (2021) Analysis of accumulation and phytotoxicity mechanism of uranium and cadmium in two sweet potato cultivars. Journal of Hazardous Materials 409: 124997

Lai J, Liu Z, Luo X (2020) A metabolomic, transcriptomic profiling, and mineral nutrient metabolism study of the phytotoxicity mechanism of uranium. Journal of Hazardous Materials 386: 121437

Laohavisit A, Shang Z, Rubio L, Cuin TA, Véry A-A, Wang A, Mortimer JC, Macpherson N, Coxon KM, Battey NH, et al (2012) Arabidopsis Annexin1 Mediates the Radical-Activated Plasma Membrane Ca2+- and K+-Permeable Conductance in Root Cells. The Plant Cell 24: 1522–1533

Lara A, Ródenas R, Andrés Z, Martínez V, Quintero FJ, Nieves-Cordones M, Botella MA, Rubio F (2020) Arabidopsis K+ transporter HAK5-mediated high-affinity root K+ uptake is regulated by protein kinases CIPK1 and CIPK9. Journal of Experimental Botany 71: 5053–5060

Laurette J, Larue C, Llorens I, Jaillard D, Jouneau P-H, Bourguignon J, Carrière M (2012a) Speciation of uranium in plants upon root accumulation and root-to-shoot translocation: A XAS and TEM study. Environmental and Experimental Botany 77: 87–95

Laurette J, Larue C, Mariet C, Brisset F, Khodja H, Bourguignon J, Carrière M (2012b) Influence of uranium speciation on its accumulation and translocation in three plant species: Oilseed rape, sunflower and wheat. Environmental and Experimental Botany 77: 96–107

Lee S, Lee EJ, Yang EJ, Lee JE, Park AR, Song WH, Park OK (2004) Proteomic Identification of Annexins, Calcium-Dependent Membrane Binding Proteins That Mediate Osmotic Stress and Abscisic Acid Signal Transduction in Arabidopsis. Plant Cell 16: 1378–1391

Lindsey BE, Rivero L, Calhoun CS, Grotewold E, Brkljacic J (2017) Standardized Method for High-throughput Sterilization of Arabidopsis Seeds. JoVE 56587

Mao D, Chen J, Tian L, Liu Z, Yang L, Tang R, Li J, Lu C, Yang Y, Shi J, et al (2014) Arabidopsis Transporter MGT6 Mediates Magnesium Uptake and Is Required for Growth under Magnesium Limitation. The Plant Cell 26: 2234–2248

Misson J, Henner P, Morello M, Floriani M, Wu T-D, Guerquin-Kern J-L, Février L (2009) Use of phosphate to avoid uranium toxicity in Arabidopsis thaliana leads to alterations of morphological and physiological responses regulated by phosphate availability. Environmental and Experimental Botany 67: 353–362

Muller DS, Houpert P, Cambar J, Hengé-Napoli M-H (2008) Role of the Sodium-Dependent Phosphate Cotransporters and Absorptive Endocytosis in the Uptake of Low Concentrations of Uranium and Its Toxicity at Higher Concentrations in LLC-PK1 Cells. Toxicological Sciences 101: 254–262

Nakagawa Y, Katagiri T, Shinozaki K, Qi Z, Tatsumi H, Furuichi T, Kishigami A, Sokabe M, Kojima I, Sato S, et al (2007) Arabidopsis plasma membrane protein crucial for Ca2+ influx and touch sensing in roots. PNAS 104: 3639–3644

Qi L, Basset C, Averseng O, Quéméneur E, Hagège A, Vidaud C (2014) Characterization of UO22+ binding to osteopontin, a highly phosphorylated protein: insights into potential mechanisms of uranyl accumulation in bones†. Metallomics 6: 166–176

Rajabi F, Jessat J, Garimella JN, Bok F, Steudtner R, Stumpf T, Sachs S (2021) Uranium(VI) toxicity in tobacco BY-2 cell suspension culture – A physiological study. Ecotoxicology and Environmental Safety 211: 111883

RStudio Team (2015) RStudio: Integrated Development for R. RStudio, Inc., Boston, MA.

Saenen E, Horemans N, Vanhoudt N, Vandenhove H, Biermans G, van Hees M, Wannijn J, Vangronsveld J, Cuypers A (2015) Oxidative stress responses induced by uranium exposure at low pH in leaves of Arabidopsis thaliana plants. Journal of Environmental Radioactivity 150: 36–43

Saenen E, Horemans N, Vanhoudt N, Vandenhove H, Biermans G, Hees MV, Wannijn J, Vangronsveld J, Cuypers A (2013) Effects of pH on uranium uptake and oxidative stress responses induced in Arabidopsis thaliana. Environmental Toxicology and Chemistry 32: 2125–2133

Sager R, Lee J-Y (2014) Plasmodesmata in integrated cell signalling: insights from development and environmental signals and stresses. Journal of Experimental Botany 65: 6337–6358

Sarthou MCM, H. Revel B, Villiers F, Alban C, Bonnot T, Gigarel O, Boisson A-M, Ravanel S, Bourguignon J (2020) Development of a metalloproteomic approach to analyse the response of Arabidopsis cells to uranium stress. Metallomics 12: 1302–1313

Serre NBC, Alban C, Bourguignon J, Ravanel S (2019) Uncovering the physiological and cellular effects of uranium on the root system of Arabidopsis thaliana. Environmental and Experimental Botany 157: 121–130

Tang R-J, Luan S (2017) Regulation of calcium and magnesium homeostasis in plants: from transporters to signaling network. Current Opinion in Plant Biology 39: 97–105

Tang R-J, Zhao F-G, Garcia VJ, Kleist TJ, Yang L, Zhang H-X, Luan S (2015) Tonoplast CBL– CIPK calcium signaling network regulates magnesium homeostasis in Arabidopsis. Proc Natl Acad Sci USA 112: 3134–3139

Tawussi F, Walther C, Gupta DK (2017) Does low uranium concentration generates phytotoxic symptoms in Pisum sativum L. in nutrient medium? Environ Sci Pollut Res 24: 22741–22751

Tewari R, Horemans N, Nauts R, Wannijn J, Van Hees M, Vandenhove H (2015) Uranium exposure induces nitric oxide and hydrogen peroxide generation in Arabidopsis thaliana. Environmental and Experimental Botany 120: 55–64

Vandenhove H (2002) European sites contaminated by residues from the ore-extracting and - processing industries. International Congress Series 1225: 307–315

Vanhoudt N, Cuypers A, Horemans N, Remans T, Opdenakker K, Smeets K, Bello DM, Havaux M, Wannijn J, Van Hees M, et al (2011a) Unraveling uranium induced oxidative stress related responses in Arabidopsis thaliana seedlings. Part II: responses in the leaves and general conclusions. Journal of Environmental Radioactivity 102: 638–645

Vanhoudt N, Horemans N, Biermans G, Saenen E, Wannijn J, Nauts R, Van Hees M, Vandenhove H (2014) Uranium affects photosynthetic parameters in Arabidopsis thaliana. Environmental and Experimental Botany 97: 22–29

Vanhoudt N, Vandenhove H, Horemans N, Bello MDM, Hees MV, Wannijn J, Carleer R, Vangronsveld J, Cuypers A (2011b) Uranium Induced Effects on Development and Mineral Nutrition of Arabidopsis Thaliana. Journal of Plant Nutrition 34: 1940–1956

Vanhoudt N, Vandenhove H, Horemans N, Remans T, Opdenakker K, Smeets K, Bello DM, Wannijn J, Van Hees M, Vangronsveld J, et al (2011c) Unraveling uranium induced oxidative stress related responses in Arabidopsis thaliana seedlings. Part I: responses in the roots. Journal of Environmental Radioactivity 102: 630–637

Vert G, Grotz N, Dédaldéchamp F, Gaymard F, Guerinot ML, Briat J-F, Curie C (2002) IRT1, an Arabidopsis Transporter Essential for Iron Uptake from the Soil and for Plant Growth. Plant Cell 14: 1223–1233

Wang J, Tergel T, Chen J, Yang J, Kang Y, Qi Z (2015) Arabidopsis transcriptional response to extracellular Ca2+ depletion involves a transient rise in cytosolic Ca2+. Journal of Integrative Plant Biology 57: 138–150

Wetterlind J, Forges ACRD, Nicoullaud B, Arrouays D (2012) Changes in uranium and thorium contents in topsoil after long-term phosphorus fertilizer application. Soil Use and Management 28: 101–107

Wheeler H, Hanchey P (1971) Pinocytosis and Membrane Dilation in Uranyl-Treated Plant Roots. Science 171: 68–71

Wilkins KA, Matthus E, Swarbreck SM, Davies JM (2016) Calcium-Mediated Abiotic Stress Signaling in Roots. Front Plant Sci 7: 1296–1296

Wu R, Fan Y, Wu Y, Zhou S, Tang S, Feng X, Tan X, Wang J, Liu L, Jin Y, et al (2020) Insights into mechanism on organic acids assisted translocation of uranium in Brassica juncea var. foliosa by EXAFS. Journal of Environmental Radioactivity 218: 106254

Yamanaka T, Nakagawa Y, Mori K, Nakano M, Imamura T, Kataoka H, Terashima A, Iida K, Kojima I, Katagiri T, et al (2010) MCA1 and MCA2 That Mediate Ca2+ Uptake Have Distinct and Overlapping Roles in Arabidopsis. PLANT PHYSIOLOGY 152: 1284–1296

Yoshimoto N, Takahashi H, Smith FW, Yamaya T, Saito K (2002) Two distinct high-affinity sulfate transporters with different inducibilities mediate uptake of sulfate in Arabidopsis roots. The Plant Journal 29: 465–473

Zhang T, Yang J, Sun Y, Kang Y, Yang J, Qi Z (2018) Calcium deprivation enhances non-selective fluid-phase endocytosis and modifies membrane lipid profiles in Arabidopsis roots. Journal of Plant Physiology 226: 22–30

